# Extended snake venomics by top-down in-source decay: Investigating the newly discovered Anatolian Meadow viper subspecies, *Vipera anatolica senliki*

**DOI:** 10.1101/773606

**Authors:** Benjamin-Florian Hempel, Maik Damm, Mrinalini, Bayram Göçmen, Mert Karış, Ayse Nalbantsoy, R. Manjunatha Kini, Roderich D. Süssmuth

**Affiliations:** Department of Chemistry, Technical University Berlin, Straße des 17. Juni 124, 10623 Berlin, Germany; Department of Biological Sciences, National University of Singapore, 14 Science Drive 4, Singapore 117543; Department of Biology, Faculty of Science, Ege University, 35100 Bornova, Izmir, Turkey; Department of Bioengineering, Faculty of Engineering, Ege University, Bornova, 35100 Izmir, Turkey; Department of Chemistry and Chemical Process Technologies, Acıgöl Vocational High School of Technical Sciences, Nevşehir Hacı Bektaş Veli University, 50300 Nevşehir, Turkey

**Keywords:** Anatolian Meadow viper, Bottom-up, In-source decay, Intact mass profiling, Venomics, Transcriptomics, Proteomics, Top-down, *Vipera anatolica senliki*, *Viperidae*

## Abstract

Herein we report on the venom proteome of *Vipera anatolica senliki,* a recently discovered and hitherto unexplored subspecies of the critically endangered Anatolian Meadow viper endemic to the Antalya Province of Turkey. Integrative venomics, including venom gland transcriptomics as well as complementary bottom-up and top-down proteomic analyses, were applied to fully characterize the venom of *V. a. senliki*. Furthermore, the classical top-down venomics approach was extended to elucidate the venom proteome by an alternative in-source decay (ISD) proteomics workflow using the reducing matrix 1,5-diaminonaphthalene (1,5-DAN). Top-down ISD proteomics allows for disulfide bond mapping as well as effective *de novo* identification of high molecular weight venom constituents, both of which are difficult to achieve by commonly established top-down approaches. Venom gland transcriptome analysis identified 42 venom transcript annotations from 13 venom toxin families. Relative quantitative snake venomics revealed snake venom metalloproteinases (svMP, 42.9%) as the most abundant protein family, followed by several less dominant toxin families. Online mass profiling and top-down venomics provide a detailed insight into the venom proteome of *V. a. senliki* and facilitates a comparative analysis of venom variability for the closely related subspecies, *V. a. anatolica*.

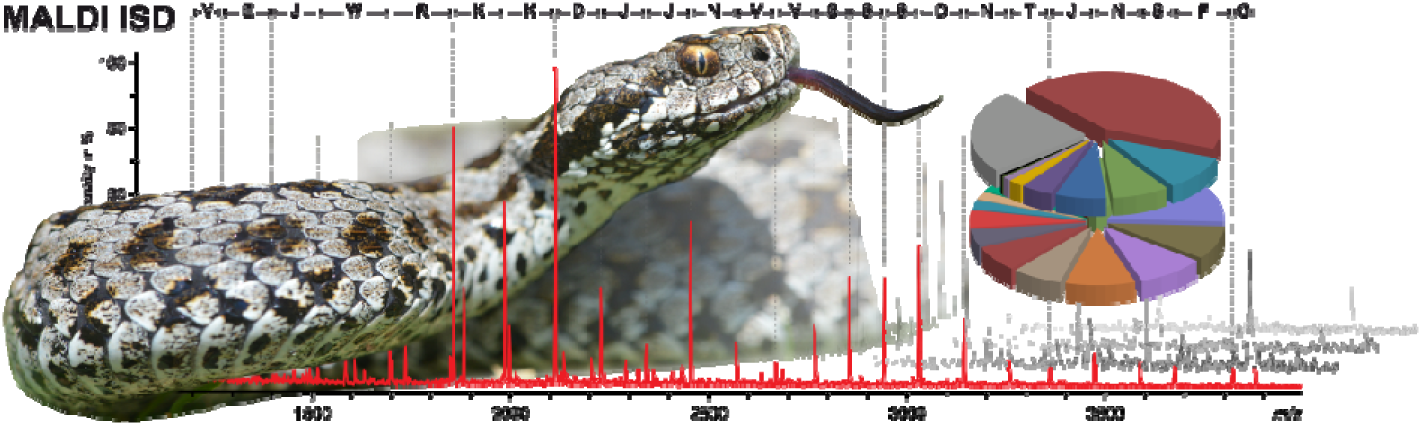
TOC Figure.

## 1. Introduction

Venoms are fascinating biological cocktails mainly composed of peptides as well as proteins and are produced by a phylogenetically broad range of organisms in terms of predation, competition or defense.^1^ The global analyses of venom for diverse members of *Serpentes* are termed as ‘snake venomics’, which usually employs an integrative approach combining proteomics, transcriptomics and occasionally genomics.^2–4^ The term was coined in 2004 by Juarez *et al.*^5^, who described for the first time the decomplexation of snake venom by a combination of liquid chromatography (LC) separation in the first dimension, followed by a one-dimensional electrophoresis (SDS-PAGE) in the second dimension.^6^ Since then, the venomics community experienced a rapid expansion into different venomic applications and a multitude of proteome studies for a wide variety of snake families have been published.^7, 8^ Today, decomplexation of snake venom proteomes can be achieved by several bottom-up protocols, combining multidimensional separation methods.^7, 9^ However, bottom-up proteomics suffers from classical proteolytic drawbacks, a step that often obscures the differentiation of toxin proteoforms and prevents the identification of post-translational modifications (PTMs).^10–13^ In this context, the recently introduced top-down protocol for snake venoms provide remedy by directly analyzing intact proteins and peptides out of crude venoms using high-resolution tandem mass spectrometry.^14–17^ However, in case of higher molecular weight compounds (>30 kDa), which are typically strongly represented in the genus of *Viperidae*, the top-down analysis only provides a partial characterization and is still challenging.^18–21^

Here, we applied a *de novo* in-source decay-driven (ISD) venomic workflow using matrix-assisted laser desorption/ionization (MALDI) as an alternative top-down approach in order to characterize high molecular mass venom constituents and to identify possible PTMs. MALDI-ISD refers, in contrast to the commonly post-source decay (PSD), to fragmentation directly in the MALDI plume during the desorption/ionization prior ion extraction.^22, 23^ The pseudo-MS/MS technique uses the hydrogen radical transfer from the 1,5-diamononaphthalene (1,5-DAN) matrix to the analyte, thus providing predominantly c- and z-type fragmentation.^24–27^ The MALDI-ISD top-down sequencing (TDS) is used to characterize protein terminal sequences from the N- or C-termini of intact proteins with the exception of roughly 10 terminally located amino acids due to matrix adducts and/or high background noise. Access to these terminal amino acids can be provided by pseudo-MS3 fragmentation, termed T^3^-sequencing, using tandem mass spectrometry (MS/MS) analysis of the previously ISD-generated fragment ions.^28, 29^ Several studies on MALDI-ISD have shown the applicability to identify high molecular mass proteoforms as well as to assign post-translational modifications (PTMs).^22, 30–35^ A special characteristic of MALDI-ISD *de novo* sequencing are short gaps in the c- or z-ion series caused by proline (Pro) residues (secondary amine), named ‘proline gap’, due to its cyclic nature. For this reason, a [M - 97]^+^ gap in the c-ion series can be interpreted as a Pro residue downstream of a preceding N-terminal amino acid.^36, 37^ In addition, it was reported that the matrix 1,5-DAN can act as a reducing agent and facilitates disulfide bond mapping, which is an accessory technique to identify snake venom families by their characteristic number of disulfide bridges.^9, 38^

A proof-of-concept study by Quinton *et al.*^39^ showed the applicability of ISD for small individual and highly purified snake peptides (<10 kDa) such as cardiotoxins and neurotoxins from the Monocled Cobra (*Naja kaouthia*). Here, we take this application one step further by characterizing the entire venom of the recently discovered Anatolian Meadow viper subspecies, *Vipera anatolica senliki*, by MALDI-ISD in combination with established snake venomic protocols for a comprehensive venom mapping. The Anatolian Meadow viper *Vipera anatolica* (*V. anatolica*) is a small, mainly insectivorous species from high altitude, stony grasslands (alpine to sub-alpine meadows). Until 2017, *V. anatolica*, was classified as a monotypic species with a narrow distribution area in Kohu Mountain, Çığlıkara Cedar Reserve, Elmali district, western Antalya province (Turkey).^40^ The venom proteome was previously described and is composed of snake venom metalloproteases (svMP), cysteine-rich secretory proteins (CRISP), phospholipases A_2_ (PLA_2_), disintegrins (DI), snake venom serine proteases (svSP), C-type lectin-like proteins (CTL), Kunitz-type protease inhibitors (KUN) and various small peptides.^18^ In a recent field study by Göçmen *et al.*^41^ a new population of the rare Anatolian meadow viper was discovered at the Senir, Serinyaka and Gelesandra Plateaus of Gündoğmuş district in eastern Antalya province, around 200 km distant from the known habitats. Subsequent, morphological and genetic analysis against known *V. anatolica* (now *V. a. anatolica*; Vaa) populations allowed a taxonomic classification and is designated *Vipera anatolica senliki* (*V. a. senliki*; Vas) as new subspecies.^41^ The combination of venom gland transcriptomics, decomplexing bottom-up proteomics, intact mass profiling and ISD top-down proteomics of *V. a. senliki* venom shed further light on the detailed venom composition of *V. a. senliki* and subspecies venom variations.

## 2. Material & Methods

### 2.1. Sampling of crude venom and venom gland dissection

Venom samples were pooled from five female and five male specimens of *V. a. senliki* from the Mühür Mountain, Senir and Gelesandra Plateaus, Gündoğmuş district (Antalya Province, Turkey) between May and June 2016. Ethical permission (Ege University Animal Experiments Ethics Committee, 2010#43) and special permission (2014#51946) for field studies from the Republic of Turkey, Ministry of Forestry and Water Affairs were obtained.

One male specimen was used for venom gland collection. Dissection was performed on a sterile surface under general anaesthesia by intramuscular ketamine injection. The venom gland lies just behind the eye and is covered by the *compressor glandulae* muscle. Primarily, this muscle was dissected to uncover the venom gland. The venom gland was gently dissected and preserved in RNA*later*™ (Thermo, Bremen, Germany) stabilization solution.

### 2.2. Venom gland transcriptomics

#### 2.2.1. RNA extraction and transcriptome sequencing

The venom gland was shipped to the National University of Singapore on dry ice and transferred immediately to -80 °C freezer for storage until RNA extraction. The sample was thawed on ice and homogenized in 400 µL RNAzol^®^ RT using a homogenizer and polypropylene pestle. An additional 600 µL of RNAzol^®^ RT was added and mixed in with the homogenate. Total RNA was extracted as per manufacturer’s protocol (in brief, DNA, proteins, and polysaccharides were eliminated and ethanol washes were performed to purify RNA). The RNA pellet was resuspended in 22 µL molecular biology grade water. Preliminary RNA quality checks and quantification was performed on NanoVue Plus. Transcriptome cDNA library preparation and sequencing were performed at NovogeneAIT. Total RNA quality and concentration was assessed using Bioanalyzer and 2 µg was used for mRNA enrichment. cDNA library was constructed using the NEBNext^®^ Ultra^TM^ Directional Library Prep Kit followed by library quality check on Bioanalyzer. 250bp Paired End (PE) sequencing was performed on 1/14^th^ of a lane on Illumina HiSeq 2500 platform.

#### 2.2.2. Transcriptome assembly

The 250bp PE raw reads were quality checked using FastQC^42^ (v0.11.4) (http://www.bioinformatics.babraham.ac.uk/projects/fastqc/) and then processed in Trimmomatic- 0.32^43^ to trim adapters and primers. Quality trimming was performed using a sliding window Phred score of Q30 for a high confidence read set. A *de novo* approach was used to assemble transcriptomes and employed two assemblers including VT-builder^44^ (v7.02), a dedicated assembler for venom genes, and the generic assembler Trinity^45, 46^ (v2.6.6). Both assemblies were then combined and de-replicated at 100% identity using CD-HIT^47, 48^ (v4.6.7).

#### 2.2.3. Venom transcriptome database

All de-replicated contigs were searched against a curated venom database downloaded from Uniprot. BLASTX was performed with an *e-value* cut off of 1e-05 in DIAMOND^49^ (v0.8.38.100). All significant hits were quality filtered using an alignment length and sequence identity of ≥60% each. Filtered sequences were aligned to venom toxin sequences in the database and then manually curated for accuracy. Venom gene expression was quantified by mapping the short reads back to the *de novo* transcriptome assembly using Salmon^50^ (v0.9.1).

### 2.3. Preparation of venom sample for top-down venomics

The crude venoms were dissolved in aqueous 1% (v/v) formic acid (FA), to a final concentration of 10 mg/mL, and centrifuged at 20,000 x *g* for 5 min to spin down insoluble content. Next, dissolved venom was mixed with 10 µL of tris(2-carboxyethyl)-phosphine (TCEP, 0.5 M) and 30 µL of citrate buffer (0.1 M, pH 3.0) to chemically reduce disulfide bonds. The reaction mixture was incubated for 30 min at 65 °C. In the following, the sample was mixed with an equal volume of 1% formic acid and centrifuged at 20,000 x *g* for 5 min. Then, 10 µL of both, reduced and non-reduced samples, were submitted for HPLC-high-resolution (HR) ESI-MS/MS measurements.

### 2.4. Top-down venomics

The top-down MS analysis was performed by dissolving crude venoms in 1% FA in ultrapure water to a final concentration of 10 mg/mL, and centrifuged at 20,000 x *g* for 5 min. The intact mass profile and top-down analysis were performed by LC-ESI-HR-MS experiments on an LTQ Orbitrap XL mass spectrometer (Thermo, Bremen, Germany) coupled to an Agilent 1200 HPLC system (Agilent, Waldbronn, Germany), using a Supelco Discovery 300 Å C18 (2 x 150 mm, 3 mm particle size) column. The flow rate was set to 0.3 mL/min and the column was eluted with a gradient of 0.1% FA in water (solution A) and 0.1% FA in acetonitrile (ACN) (solution B): 5% B for 1 min, followed by 5–40% B for 89 min, and 40–70% for 20 min. Finally, the column was washed with 70% B for 10 min and re-equilibrated at 5% B for 10 min. ESI settings were 45 L/min sheath gas; 5 L/min auxiliary gas; spray voltage, 4.5 kV; capillary voltage, 46 V; tube lens voltage, 155 V; and capillary temperature, 275 °C. MS/MS spectra were obtained in data-dependent acquisition (DDA) mode. FTMS measurements were performed with 1 μ scans and 500 ms maximal fill time. AGC targets were set to 10^6^ for full scans and to 1 × 10^5^ for MS/MS scans and the survey scan as well as both data dependent MS/MS scans were performed with a mass resolution (R) of 100,000 (at *m*/*z* 400). For MS/MS the top three most abundant ions of the survey scan with known charge were selected. Normalized CID energy was set to 30% for the first, and 35% for the second MS/MS event of each duty cycle. The default charge state was set to z = 9, and the activation time to 30 ms. Additional HCD experiments were performed with 35% normalized collision energy, 30 ms activation time and z = 9 default charge state. The mass window for precursor ion selection was set to 2 or 4 *m*/*z*. A window of 3 *m*/*z* was set for dynamic exclusion of up to 50 precursor ions with a repeat of 1 within 10 s for the next 20 s. The intact mass profile was inspected with the Xcalibur Qual Browser (Thermo Xcalibur 2.2 SP1.48) and deconvolution of isotopically resolved spectra was carried out by using the XTRACT algorithm of Xcalibur Qual Browser. Protein assignment was done by comparison to the retention time of the HPLC run.

### 2.5. Chromatographic separation of venom sample

The lyophilized crude venom (2 x 1 mg) was dissolved to a final concentration of 20 mg/mL in aqueous 3% (v/v) ACN with 1% (v/v) FA and centrifuged at 16,000 x *g* for 5 min to spin down insoluble content. The supernatant was loaded onto a semi-preparative reversed-phase HPLC (RP-HPLC) with a Supelco Discovery BIO wide Pore C18-3 column (4.6 x 150 mm, 3 µm particle size) using an Agilent 1260 High pressure Gradient System (Agilent, Waldbronn, Germany). The column was operated with a flow rate of 1 mL/min and performed ultrapure water with 0.1% (v/v) FA (buffer A) and ACN with 0.1% (v/v) FA (buffer B). We used a standard separation gradient with solution A and solution B, starting isocratically (5% B) for 5 min, followed by linear gradients of 5-40% B for 95 min and 40-70% for 20 min, then 70% B for 10 min, and finally re-equilibration at 5% B for 10 min. Detection was performed at λ = 214 nm using a diode array detector (DAD). After the chromatographic separation fractions were vacuum-dried and used for bottom-up analysis and in source decay analysis.

### 2.6. Bottom-up characterization and quantification of the venom proteome

The dried fractions were submitted to SDS-PAGE (12% polyacrylamide). Afterwards the Coomassie-stained band were excised and via in-gel trypsin digestion reduced with fresh dithiothreitol (100 mM DTT in 100 mM ammonium hydrogen carbonate, pH 8.3, for 30 min at 56 °C) and alkylated with iodoacetamide (55 mM IAC in 100 mM ammonium hydrogen carbonate, pH 8.3, for 30 min at 25 °C in the dark). An in-gel trypsin (Thermo, Rockfeld, IL, USA) digestion was performed (6.7 ng/µL in 10 mM ammonium hydrogencarbonate with 10% (v/v) ACN, pH 8.3, for 18 h at 37 °C, 0.27 µg/band). The peptides were extracted with 100 µL aqueous 30% (v/v) ACN just as 5% (v/v) FA for 15 min at 37 °C. The supernatant was vacuum dried (Thermo SpeedVac, Bremen, Germany), redissolved in 20 µL aqueous 3% (v/v) ACN with 1% (v/v) FA and submitted to LC-MS/MS analysis.

The bottom-up analysis were performed with an Orbitrap XL mass spectrometer (Thermo, Bremen, Germany) via an Agilent 1260 HPLC system (Agilent Technologies, Waldbronn, Germany) using a reversed-phase Grace Vydac 218MSC18 (2.1 x 150 mm, 5 µm particle size) column. The pre-chromatographic separation was performed with the following settings: After an isocratic equilibration (5% B) for 1 min, the peptides were eluted with a linear gradient of 5-40% B for 10 min, 40-99% B for 3 min, washed with 99% B for 3 min and re-equilibrated in 5% B for 3 min.

LC-MS/MS data files (.raw) were converted to mascot generic format (.mgf) files via MSConvert GUI of the ProteoWizard package (http://proteowizard.sourceforge.net; v3.0.10577) and annotated by DeNovo GUI^51^ (v1.15.11) with the following settings: fixed modifications: carbamidomethyl cysteine (+57.02 Da); variable modifications: acetylation of lysine (+42.01 Da), and phosphorylation of serine and threonine (+79.97 Da). The deduced peptide sequences were squared against a non-redundant protein NCBI database of *Viperidae* (taxid: 8689) using the basic local alignment tool (BLAST)^52^ (http://blast.ncbi.nlm.nih.gov).

For peptide spectrum matching, the SearchGUI^53^ (v3.3.11) software tool was used with XTandem! as the search engine. The MS/MS spectra were searched against the non-redundant *Viperidae* protein NCBI (taxid: 8689, 17th Dec 2018, 1310 sequences), our custom toxin sequence database from *V. a. senliki* (derived from the translation of our venom gland transcriptomics database; 42 toxin sequences) and a set of proteins found as common contaminants (**c**ommon **R**epository of **A**dventitious **P**roteins, cRAP, 116 sequences), containing in total 1,468 sequences. Precursor mass accuracy was set to 10 ppm and 0.5 Da for the MS^2^ level. The alkylation of Cys was set as a fixed modification and acetylation of the N-terminus and Lys were allowed as variable modifications. A false discovery rate (FDR) was estimated through a target-decoy approach and a cut-off of 1% was applied.

### 2.7. In-source decay

The top-down MS analysis by in source decay was performed by redissolving vacuum-dried HPLC fractions (2 x 1 mg) in ultrapure water. The redissolved fractions were mixed with the reducing matrix 1,5-diaminonaphthalene (1,5-DAN, 20 mg/mL in 50:50 [v/v] acetonitrile (TA50) and 0.1% FA in water) or 2,5-dihydroxybenzoic acid (2,5-DHB, 20 mg/mL in TA30) in a ratio of 1:1, then gently mixed and subsequently spotted on a polished ground steel plate (Bruker, Bremen, Germany). The MALDI-ISD measurements were performed on a Bruker ultraflex TOF/TOF device (Bruker, Bremen, Germany). Instrument calibration was performed using a peptide mix (Peptide Calibration Standard from Bruker Daltonics, *m/z* range: 700-3200). The laser power was set to ∼50% for experiments with 2,5-DHB and increased to ∼70% for experiments using 1,5-DAN. The resulting raw files were analyzed using the Bruker BioTools software (version 2.3). Peak picking was performed with the following settings: minimal mass difference between peaks of 0.3 Da, threshold (S/N) of 1, quality factor threshold of 20 and maximum number of peaks to 1000. For the *de novo* annotation option, the ISD-TOF extended ion list was selected, including a, c, y and z+2 ions. The following fixed modification was set for top-down analysis: reduced disulfide bridges (+2 Da). Resulting peptide sequences were searched against the non-redundant *Viperidae* protein NCBI (taxid: 8689, 17th Dec 2018, 1310 sequences), our custom *V. a. senliki* toxin sequence database (translated from our venom gland transcriptomic analysis; 42 toxin sequences) and a set of proteins found as common contaminants (cRAP, 116 sequences), containing in total 1,468 sequences.

### 2.8. Data accessibility

All transcriptome data, including sample information, RNAseq short reads, and the *de novo* transcriptome assembly, have been deposited in the NCBI database (http://www.ncbi.nlm.nih.gov/sra) with the BioProject identifier PRJNA560445. Mass spectrometry proteomics data (.mgf, .raw, .fid and output files) have been deposited with the ProteomeXchange Consortium^54^ (http://proteomecentral.proteomexchange.org) via the MassIVE partner repository under project name “Extended snake venomics on the newly discovered subspecies *Vipera anatolica senliki*” with data set identifier PXD014805.

### 2.9. Relative toxin quantification

The quantification of venom composition is based on the RP-HPLC fraction integration (UV_214nm_) in comparison to the total integral of all analyzed fractions based on the protocol of Juarez *et al.*^5^. In case of co-eluting toxins components, observable by SDS Coomassie staining, the ratio of optical intensities and densities was respectively used for emphasis of fraction integral portion.

## 3. Results & Discussion

### 3.1. The venom of *Vipera anatolica senliki*

Herein we describe the detailed venom composition of the recently discovered subspecies of the Anatolian Meadow viper, *Vipera anatolica senliki,* by a combination of decomplexing bottom-up and top-down proteomic approaches as well as venom gland transcriptomics. RNA sequencing of the venom gland was followed by a *de novo* approach to assemble the venom transcriptome for *V. a. senliki*. Further, we used established protocols for a comprehensive characterization and relative quantification of the venom as a basis to subsequently describe MALDI-ISD experiments and subspecies level comparison of venom variation. The initial analysis of *V. a. senliki* venom was performed by a one-dimensional RP-HPLC separation, followed by a subsequent SDS-PAGE separation and in-gel trypsin digestion as well as a direct online intact mass analysis by ESI-HR-MS. All *de novo* and peptide sequence matches from our transcriptomic and proteomic approaches are listed in more detail in Table 1 and Supplemental Tables 1 & 2.

**Table 1.**
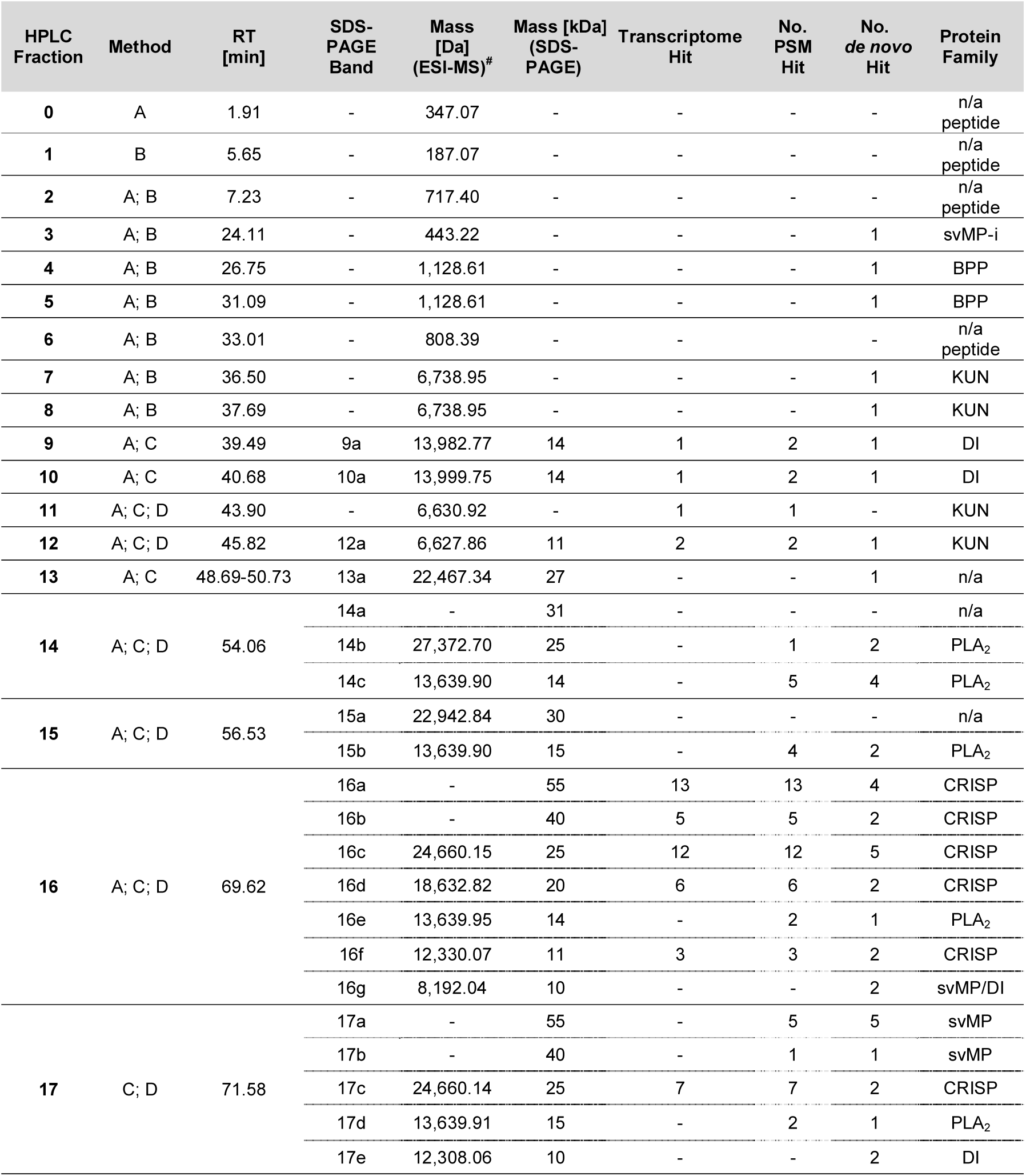
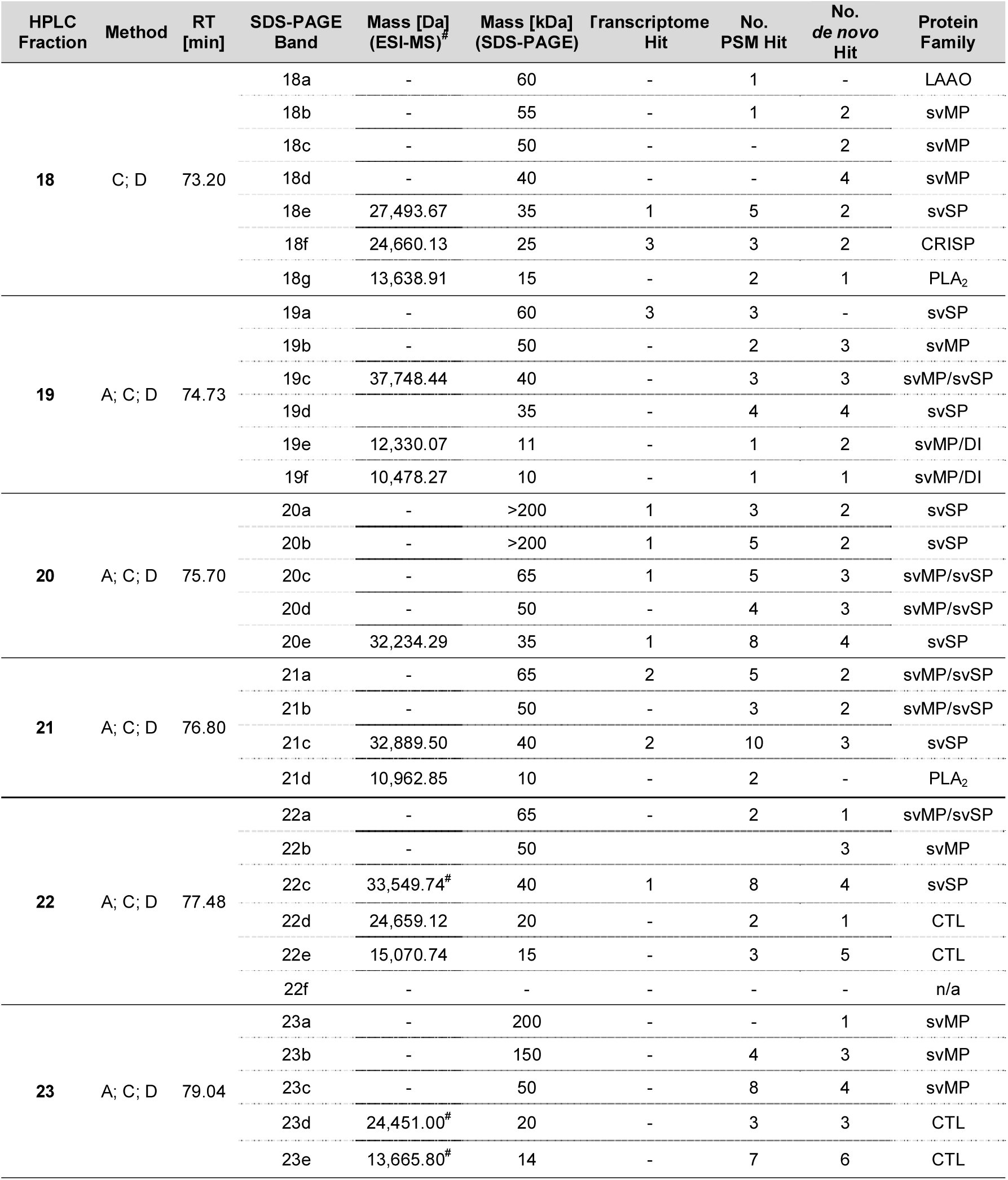
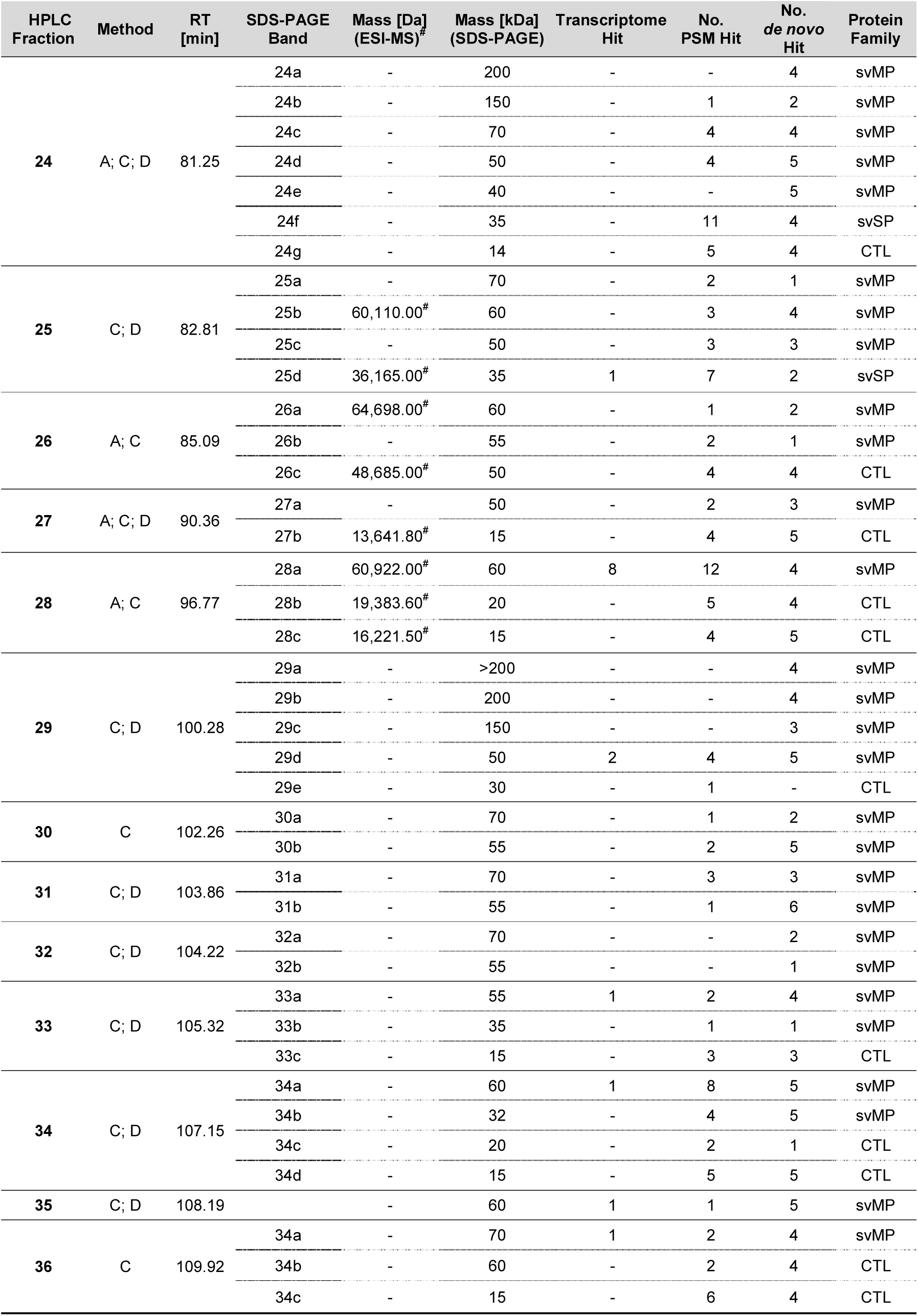
Venom proteins and peptides identified from Vipera anatolica senliki. Assignment of venomic components was performed by crude venom intact mass profiling (IMP, method A), IMP of a single RP-HPLC fraction with low molecular mass (method B), bottom-up (BU, method C) and in-source decay annotation (ISD, method D). Fraction numbers are based on the RP-HPLC chromatogram (**Figure 1**). Annotation was performed *de novo* and by peptide spectrum matching from in-gel digested protein bands (**SI-Figure 1**). Identification was carried out against a non-redundant *Viperidae* protein database (taxid: 8689), our custom transcriptome database and a set of proteins found as common contaminants (cRAP). SDS-PAGE and intact mass profile analysis provided the average molecular weight. For IMP only most abundant mass is listed (all masses in **SI-Table 1**). IMP performed by charge-state deconvolution was carried out with MagicTransformer (MagTran) and is marked by #.

#### 3.2.1 Venom gland transcriptome

Our transcriptome assembly of cDNA from the venom gland of *V. a. senliki* resulted in a total of 42 contigs that belonged to 13 venom toxin families. Of these, 14 were full length toxin sequences, whereas 28 sequences were partially assembled toxins. Full length toxins were represented by svSP, KUN, Nerve Growth Factor (NGF), Vascular Endothelial Growth Factor (VEGF), Venom Factor (VF), Waprin (WAP), Cystatin (CYS), Ryncolin (RYN), Neuropeptide-Y (NPY). While alternate isoforms from some of these families were also represented as partially assembled contigs, additional families such as CRISP, svMP, 5’-Nucleotidase (5NUC), and Glutaminyl Peptide Cyclotransferase (GPC) were wholly represented by partial contigs. The full set of venom toxins retrieved from the transcriptome and the breakdown of number of full and partial isoforms in each toxin family are shown in Figure 1C and Supplemental Table 1.

**Figure 1:**
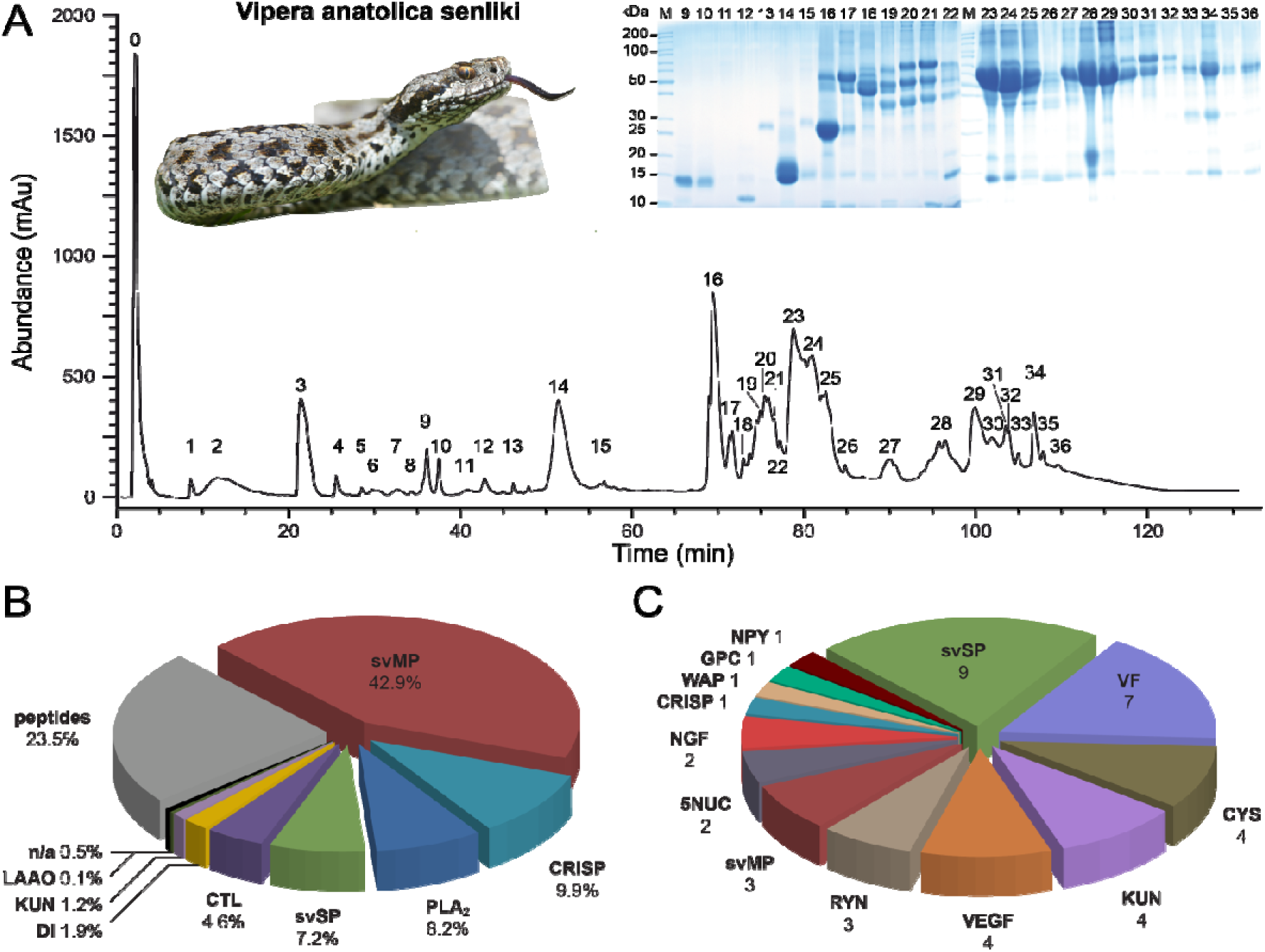
Snake venomics and venom gland transcriptomics of *Vipera anatolica senliki*. (**A**) *V. a. senliki* venom fractions of the RP-HPLC fractionation, with UV214nm detection signal, were subsequently subjected to 1D-SDS-PAGE, followed by tryptic digests of the most abundant bands. PAGE line nomenclature based on HPLC fractions. A detailed band nomenclature is shown in SI-Figure 1. The relative quantitative occurrence (**B**) and numbers of different transcripts (**C**) of the toxin families by integrative snake venomics are represented by a pie charts, with the following families: C-type lectin-like proteins (CTL), Cystatin (CYS), disintegrin (DI), glutaminyl peptide cyclotransferase (GPC), Kunitz-type protease inhibitors (KUN), L-amino acid oxidase (LAAO), not annotated (n/a), nerve growth factor (NGF), neuropeptide-Y (NPY), 5’-nucleotidase (5NUC), phospholipase A_2_ (PLA_2_), Ryncolin (RYN), snake venom metalloproteinases (svMP), snake venom serine proteases (svSP), vascular endothelial growth factor (VEGF), venom factor (VF), Waprin (WAP).

We performed an analysis of toxin gene expression in the venom gland by mapping quality filtered RNAseq reads back to the assembled venom gland transcriptome. We found that the most highly expressed toxins were CYS (42%) and KUN (34%), followed by svSP (7%), VF and Ryncolin (6% each), VEGF (2%) and WAP (1%). Other families such as svMP, CRISP, 5NUC, NGF showed negligible expression (<1%), or as in the case of GPC and NPY, no quantifiable expression was found. Interestingly, we did not find expression or even partially assembled contigs for PLA_2_. The toxin gene expression in *V. a. senliki* is atypical compared to the gene expression patterns generally reported for *Viperinae,* where PLA_2_, svMP, svSP, CTL, and CRISP have been found to dominate.^8, 55^ Prior to venom gland collection for transcriptomic studies, venom is usually extracted a few days in advance to start the gene expression cascade. Venom gene expression and protein re-synthesis is known to peak between 3-7 days after venom extraction.^56^ In the case of *V. a. senliki* however, venom extraction was performed 10-14 days in advance of gland dissection under anesthesia. Our results show that high expression of CYS (422,278 TPM) likely followed the peak expression of major venom components such as svSP, svMP, or PLA_2_. This is further supported by results from venom proteomics analysis where svMP, CRISP, and PLA_2_ were readily found to be the most abundantly represented toxin families which is in line with the typical venom composition of *Viperinae*.^8, 55^ The CYS family is thought to play a crucial role in preserving toxin potency by acting as strong inhibitors of cysteine proteases that can degrade other venom proteins.^57^ Hyper-expression of CYS, as found in our study, provides an interesting snapshot of the CYS expression pattern in the venom gland assigned to post peak venom synthesis.

#### 3.2.1. Bottom-up venomics

The decomplexation of the *V. a. senliki* venom by RP-HPLC shows 36 characteristic fractions (F), which were subsequently fractionated in the second dimension by SDS-PAGE and revealed the presence of 95 separate protein bands (**Figure 1A**). The resulting sequence tags, after in-gel trypsin digestion, were searched against the NCBI non-redundant viperid protein database using BLASTP. The 265 *de novo* annotated sequence tags resulted in the identification of several proteins covering seven toxin families (**SI-Table 1**). The most abundant protein family is represented by svMP, followed by CRISP, PLA_2_, CTL, svSP, and DI, whereas a small percentage (∼1%) could not be annotated (n/a). Another abundant part of the venom is formed by peptides, which we further investigated by the direct measurement of the nine first fractions (F 0-8) and the IMP analysis. *De novo* sequence assignment of the MS/MS spectra of small peptides (F 0-8) resulted in the identification of a snake venom metalloproteinase inhibitor (svMP-i) pEKW and a bradykinin potentiating peptide (BPP) (**SI-Table 1** and **SI-Figure 3**).

The re-analysis of the MS/MS data from the tryptic peptides was performed by peptide spectrum matching (PSM), using the assembled custom transcriptome database, the NCBI *Viperidae* protein database and a list of commonly found contaminants (cRAP). This resulted in 303 peptide matches and nine major toxin families in total. This output is a slight improvement of annotated spectra in comparison to the *de novo* annotation (**Table 1**). The vast majority of previously annotated *de novo* peptides could be confirmed by PSM analysis, but also revealed, in contrast to the *de novo* sequencing, the presence of a KUN and an L-amino acid oxidase (LAAO). Furthermore, peptide matching analysis corrected a previous false annotation from *de novo* sequencing of two serine/threonine-protein kinases and a predicted zinc-finger protein. Indeed, *de novo* sequencing is an error-prone process complicated by uneven fragmentation patterns due to missing fragmentation, limited algorithm accuracy for various interferences, homeometric peptides, low resolution, and the lack of a species-specific protein database.^58–60^ Therefore, the corresponding spectral peptide matches showed the presence of svSP and CTL instead. Overall, our PSM showed a modest improvement, but most of the obtained peptide matches were from viperid sequences in the NCBI database, with only 82 protein IDs sourced from transcriptome derived sequences. The peptide matches mainly represent hits against the transcriptome database for the toxin families svMP, svSP and CRISP. In some cases, where no hits were obtained by peptide spectrum matching, *de novo* sequence tags were resorted to classify the protein. The relative quantification showed svMP (42.9%) as most abundant toxin family, followed by CRISP (9.9%), PLA_2_ (8.2%), svSP (7.2%), CTL (4.6%), DI (1.9%), KUN (1.2%), and LAAO (0.1%). In the small molecular mass range (< 2 kDa) the following peptides were found: svMP-i (5.9%), bradykinin potentiating peptides (BPP; 0.6%) and unknown peptides (17.0%) of the overall venom composition (**Figure 1B**).

To bypass the problem of false or non-annotated peptide matches, a combination of accurate transcriptome assembly or even an alternative, more sensitive bottom-up method, like shotgun proteomics, could be useful for a targeted identification. However, shotgun proteomics suffers from a less quantitative breakdown of snake venom composition due to an inefficient differentiation of the numerous toxin isoforms present in crude venoms. Ultimately, employing top-down proteomics combined with a front-end chromatographic separation is a suitable alternative to uncover existing proteoforms, PTMs and to identify toxin components by tandem MS or disulfide bond mapping.^11, 61^

#### 3.2.2. Top-down venomics

The direct online intact mass profiling by ESI-HR-MS of *V. a. senliki* venom coupled to a front-end LC-based decomplexation resulted in a TIC profile (**SI-Figure 2**) comparable to the previous chromatographic separation (fraction nomenclature is based on Figure 1). The initial measurement of native venom generated an overview of 79 intact venom components, including low abundant compounds and small peptides. The initial fractions (F 0-8) exclusively contain smaller peptide components (>7 kDa), like svMP-i and BPP as well as a KUN proteoform (F 7 and 8). The aforementioned svMP-i (pEKW) in F 3 and BPP in F 4 and F 5 are detectable, but less prominent compared to the semi-preparative separation, which can be explained by low ionization yields of the peptides by electrospray ionization (**SI-Figure 2**). The following fractions (F 9-15) are toxin families in the molecular mass range of 10-30 kDa composed of two heterodimeric disintegrin proteoforms, two KUN proteoforms and one PLA_2_. Subsequent fractions (F 16-36) show intact masses across the entire molecular mass range from different venom protein families (10-70 kDa), like svMP, CRISP, LAAO, svSP or CTL with various proteoforms (**SI-Table 1**).

In addition to the native mass profiling, crude venom was further investigated by chemical reduction using the reagent TCEP. The comparison of native and chemically reduced venom components allows for the identification by inter- and intramolecular disulfide bonds as an important characteristic for several toxin families.^9^ A closer look to the native and reduced venom components showed the identification of three different venom protein families (**Figure 2**). The intact mass for fraction 7 in the native venom showed two distinct molecular masses (6,738.0 Da and 7,280.2 Da) in the typical range for Kunitz-type inhibitors (**Figure 2A-B**, blue). However, to completely confirm the suspected protein family, we compared it to the reduced venom profile and found the corresponding molecular masses (6,744.0 Da and 7,286.3 Da) that were shifted by Δ6.0 Da (**Figure 2A-B**, red). The individual mass shifts indicate the presence of three disulfide bridges, which is characteristic for Kunitz-type inhibitors in snake venoms.^62, 63^ In addition to the previous *de novo* annotation and PSM, we were able to identify fraction 14 as a single basic PLA_2_ by disulfide bond mapping (**Table 1**). The comparison of native (13,639.9 Da) and reduced (13,654.0 Da) mass spectra showed a mass shift of Δ14.1 Da, that correlates to seven disulfide bridges and is a clear characteristic for snake venom PLA_2_ (**Figure 2C**).^64, 65^ We were able to further characterize the three dimeric DI proteoforms (13,982.8 Da, 13,999.8 Da and 14,001.8 Da) present in fractions 9 and 10 composed of four different subunits (6,979.0 Da, 6,998.0 Da, 7,022.0 Da and 7,023.9 Da) showing two inter-subunit (Δ2.01 Da) and twice four intra-subunit (Δ8.1 Da) disulfide bridges (**Figure 2D-I**).^66, 67^ Moreover, we were able to identify small peptides, like the snake venom metalloproteinase inhibitor (svMP-i) pEKW by tandem MS (**SI-Figure 3**).

**Figure 2:**
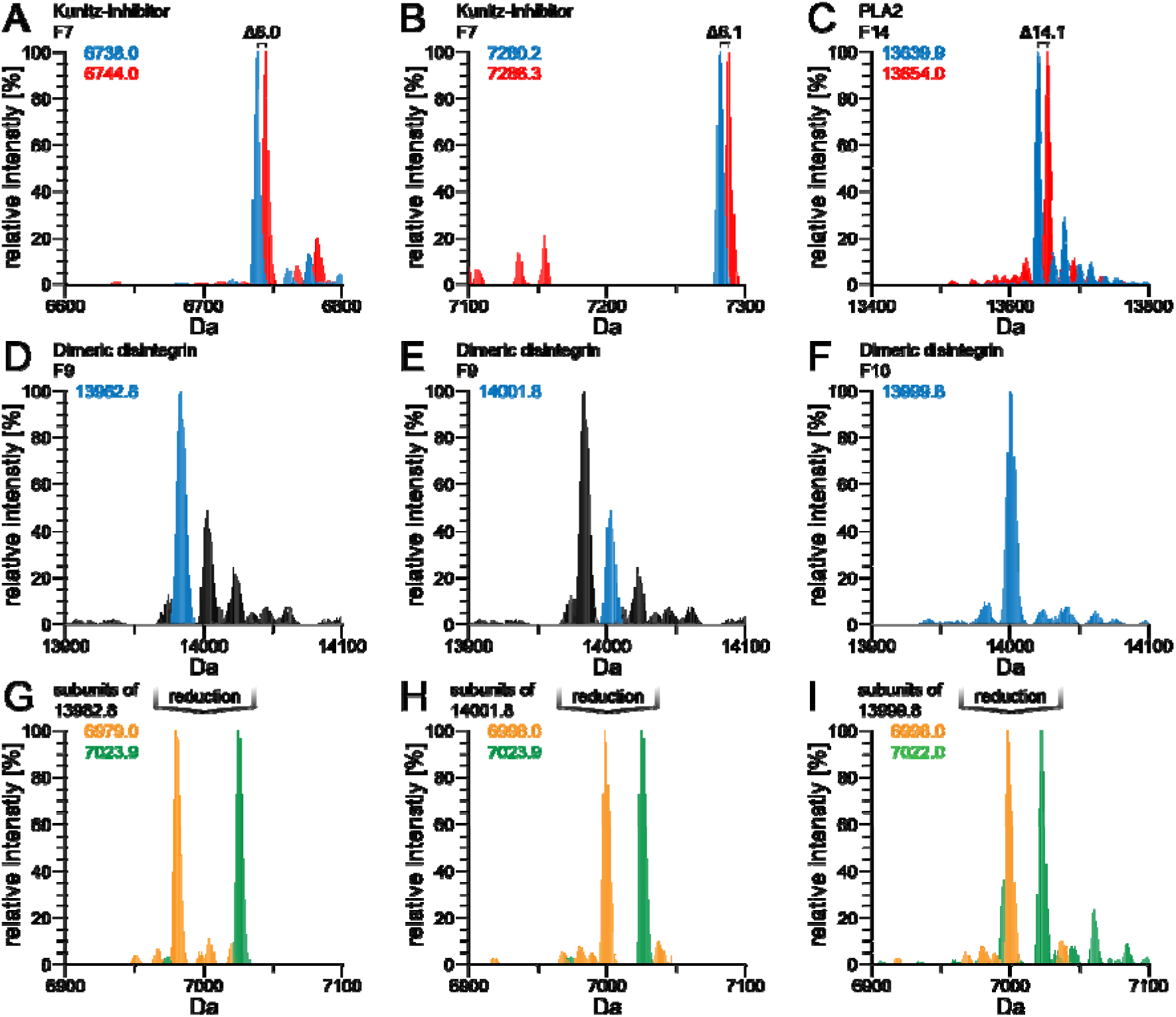
Intact mass profiling and disulfide bond mapping of selected toxin components from *Vipera anatolica senliki*. The deconvoluted native and reduced venom profiles show typical mass shifts in several fractions for three different venom protein families. (**A**, **B**) Venom fraction F7 includes two Kunitz-type serine protease (KUN) inhibitors with three (Δ6 Da) and (**C**) F14 shows a single phospholipase A2 (PLA2) with seven disulfide (S-S) bonds (Δ14 Da). (**D**, **E**, **F**) Fractions 9 and 10 include in total the three main dimeric disintegrins (DI) with two inter-subunit S-S and four intra-subunit S-S bonds each (in total Δ20 Da), shown by reduced venom (**G**, **H**, **F**).

However, the top-down venomic approach applied here suffers from distinct drawbacks of insufficient ionization by denaturing electrospray ionization and poor isotope resolution for a number of main toxin components, such as high molecular weight svMPs, that form a substantial part of viperid venoms. For this reason, we established an alternative top-down method, which offers the same advantages without the limitation of molecular weight that is ideal for venom protein families of viperid species.

### 3.3. Top-down by in-source decay

A possible way to overcome the above mentioned difficulties is the application of MALDI coupled to a time-of-flight (TOF) mass analyzer. MALDI-TOF is a valuable tool for intact mass analysis of proteins in a higher molecular mass range. In combination with 1,5-DAN as a matrix compound, which allows for a partial reduction of disulfide bonds and significantly enhanced yields of ISD fragmentation, MALDI-TOF becomes a fast and efficient method for top-down venom sequencing. Here, we go beyond a previous proof-of-concept study by Quentin *et al.*^39^ and analyze an entire snake venom to particularly identify high molecular venom components.

In order to examine the potential of ISD-derived top-down sequencing, we first fractionated the venom of *V. a. senliki* similar to the initial HPLC separation of our bottom-up venom analysis. The venom fractions were either prepared with the reductive 1,5-DAN matrix or non-reductive DHB in a ratio 1:1 and subsequently spotted on the MALDI target to allow for crystallization. The direct comparison of reduced and non-reduced spectra facilitates a classification of venom protein families by disulfide bond mapping, as described for the aforementioned top-down venomic approach. The non-reduced intact mass signal of fraction 11 shows a mass of 6,630.1 Da, which is an indicator for a Kunitz-type serine protease inhibitor (**Figure 3A**).^68, 69^ The comparison with the reduced intact mass (6,636.2 Da) reveals a mass shift of Δ6.1 Da (corresponding to three disulfide bridges), which confirms the Kunitz-type architecture. Furthermore we were able to identify several PLA_2_ proteoforms in fractions 14 and 15 by disulfide bond mapping (**Figure 3B**). The most abundant native intact masses of fractions 14 and 15 were 13,637.4 Da or 13645.6 Da, respectively, and thus in the molecular mass range of PLA_2_.^70, 71^ The corresponding reduced masses of fractions 14 and 15 (13,652.5 Da and 13,660.7 Da) show a mass shift of ∼Δ14 Da, which corresponds to seven disulfide bridges, a typical feature of PLA_2_’s. Further assignments of venom components by disulfide bond mapping were not possible due to low signal intensity of intact protein masses. The low signal intensity of the reduced intact masses (signal loss: ∼10^3^ in abundance) makes an assignment by disulfide bond mapping more difficult particularly for proteins present in low amounts.

**Figure 3:**
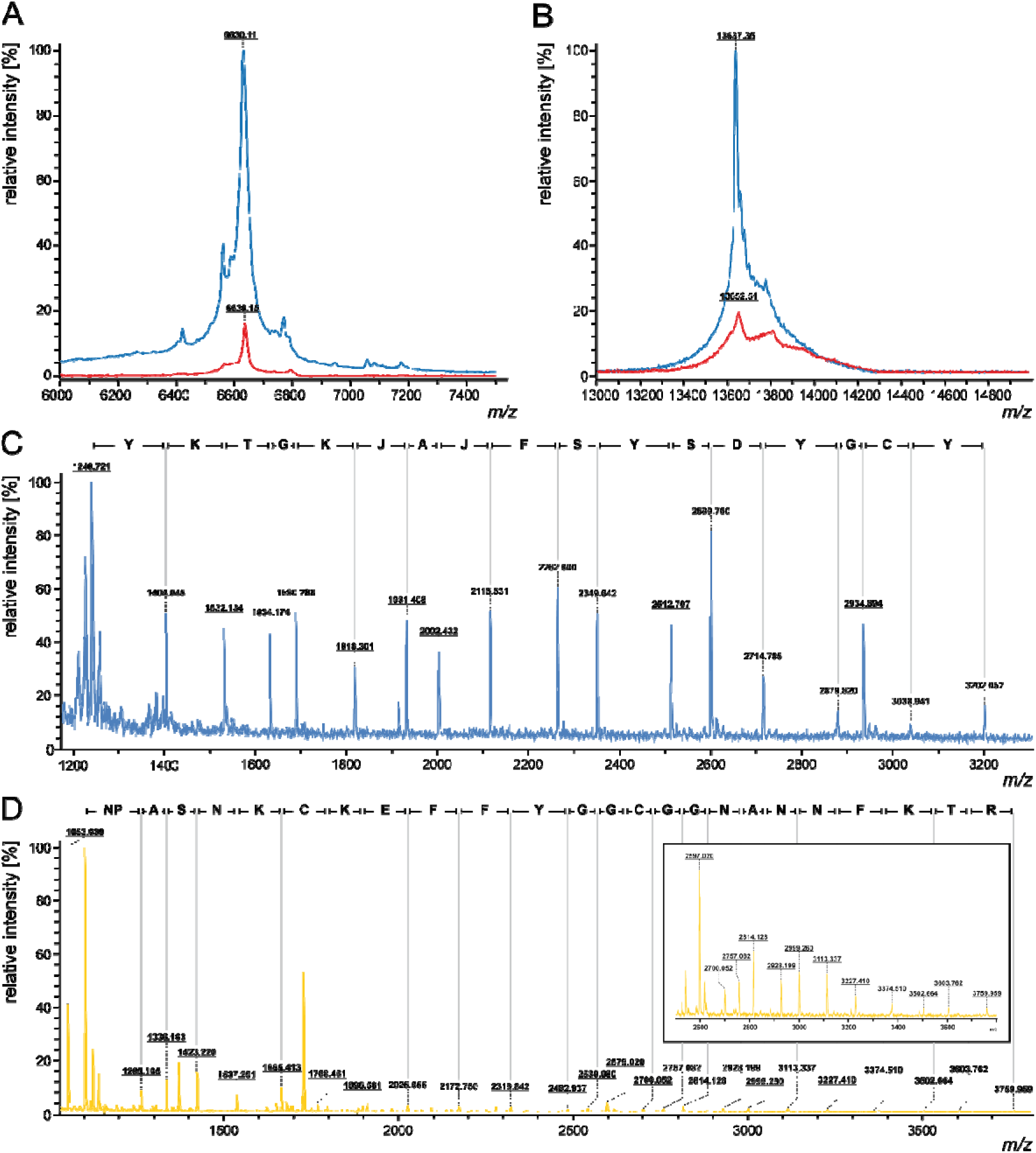
MALDI top-down sequencing by in-source decay and disulfide bond mapping of venom components from *Vipera a. senliki*. The transfer of hydrogen radicals induced by 1,5-diaminonaphthalene (1,5-DAN) matrix and the following peptide backbone cleavage allow MALDI-TOF top-down sequencing of high molecular mass toxin components by c and z+2 fragment ions. (**A**) Comparison of native (DHB) and reduced (DAN) intact mass spectra (F 11) allow identification of 3 disulfide bridges, characteristic for Kunitz-type serine protease (KUN) inhibitor. (**B**) Comparison of native (DHB) and reduced (1,5-DAN) intact mass spectra (F 14/15) allow identification of 7 disulfide bridges, characteristic for phospholipase A_2_ (PLA_2_). (**C**) Identification of a PLA_2_ proteoform by N-terminal sequence (F 14/15). (**D**) Identification of a KUN inhibitor by N-terminal sequence (F 12). No distinction can be made between leucine and isoleucine (J = Leu or Ile).

The effect contributing to much lower signal intensity is caused by hydrogen transfer and the following radical-induced cleavage of the peptide backbone. However, this is a desired process for venom proteome annotation by sequence alignment.^26, 72^ The peptide backbone cleavage caused by transfer of hydrogen radicals allows for MALDI-TOF top-down sequencing predominantly from the N-terminal part of several toxin components (**Figure 3 & 4, Table 1** and **SI-Figure 4**). The mass range below 1000 Da was only partially considered for *de novo* sequencing due to intense matrix background.^23^ Nevertheless, MALDI top-down sequencing (TDS) provides a straightforward protein sequence analysis approach for small but also high molecular mass toxin components.

In our analysis we obtained various specific c*_n_*-fragment ion series in the reflectron mode by ISD fragmentation (reISD). The aforementioned protein constituents, like KUN and PLA_2_ (F 11 & 14/15), identified by the number of disulfide bonds as well as peptide spectrum mapping, could be additionally confirmed by ISD sequence tags. The identity of the PLA_2_ (F 14/15) was underpinned by a 17-mer N-terminal sequence (YKTGKJAJFSYSDYGCY), which showed a high correlation with the PLA_2_ ammodytin I2(D) isoform (NCBI: CAE47222.1) (**Figure 3C**). Additionally, also the Kunitz-type protease inhibitor (F 12) could be assigned by an N-terminal 25-mer peptide sequence (NPASNKCKEFFYGGCGGNANNFKTR), which resulted in a specific hit to our transcriptome database (DN44715_c0_g1_i1_len_508) (**Figure 3D**). In this context, it should be mentioned that the matching transcriptome sequence is a precursor protein composed of an N-terminal signal peptide and a protein core sequence, which is post-translationally processed to the mature KUN by cleavage of the signal peptide that was finally annotated. This example shows the necessity of proteomic examination in integrative venomics expanded by the alternative ISD-driven method, which displays terminal sequences of mature proteins.

In addition to the *de novo* sequencing of venom components and their assignment to our in-house database, we confirmed the previously identified CRISP proteoform (F 16) by a 23-mer peptide fragment (KPEJQNEJJDJHNSJRRSVNPTA) to an entry of our species-specific transcriptome (DN8323_c0_g1_i1_len_755) in the N-terminal region (**Figure 4A**). Furthermore, we verified a svMP proteoform (F 17) by detection of a 22-mer peptide sequence (VEJWRKKDJJNVVSSSDNTJNS) with a high homology to a metalloproteinase (NCBI: ADI47725.1) from *Echis carinatus sochureki* (**Figure 4B**). In addition, we found a short N-terminal sequence of the CRISP proteoform, identified in fraction 16 (**SI-Figure 4D**). In fraction 29 we annotated several svMP proteoforms, which were confirmed by three 15/16-mer sequences (EJVJVVDNVMFX_1_KYX_2_ with X_1_=K/R and X_2_=K/(NG)), which are existing proteoforms in the public NCBI *Viperidae* proteome database. In addition, we identified the first annotation sequence as a modified proteoform to one of our database entries (DN2248_c0_g1_i1_len_747) (**Figure 4C**). The top-down ISD sequencing of svMP (F 31/32) matches in part our transcriptome data (DN2248_c0_g1_i1_len_747) and led to the identification of two different proteoforms with a 12-mer (YVEJVJTVDHRM) and 14-mer (JVJVVDNVMFKKYK) peptide in the N-terminal region (**Figure 4D**). These examples show the advantage of ISD top-down sequencing for the identification of related proteoforms. In addition to these examples, we confirmed several other proteoforms from various snake venom constituents by N-terminal peptide fragments of different length (**Table 1**, **SI-Figure 4**).

**Figure 4:**
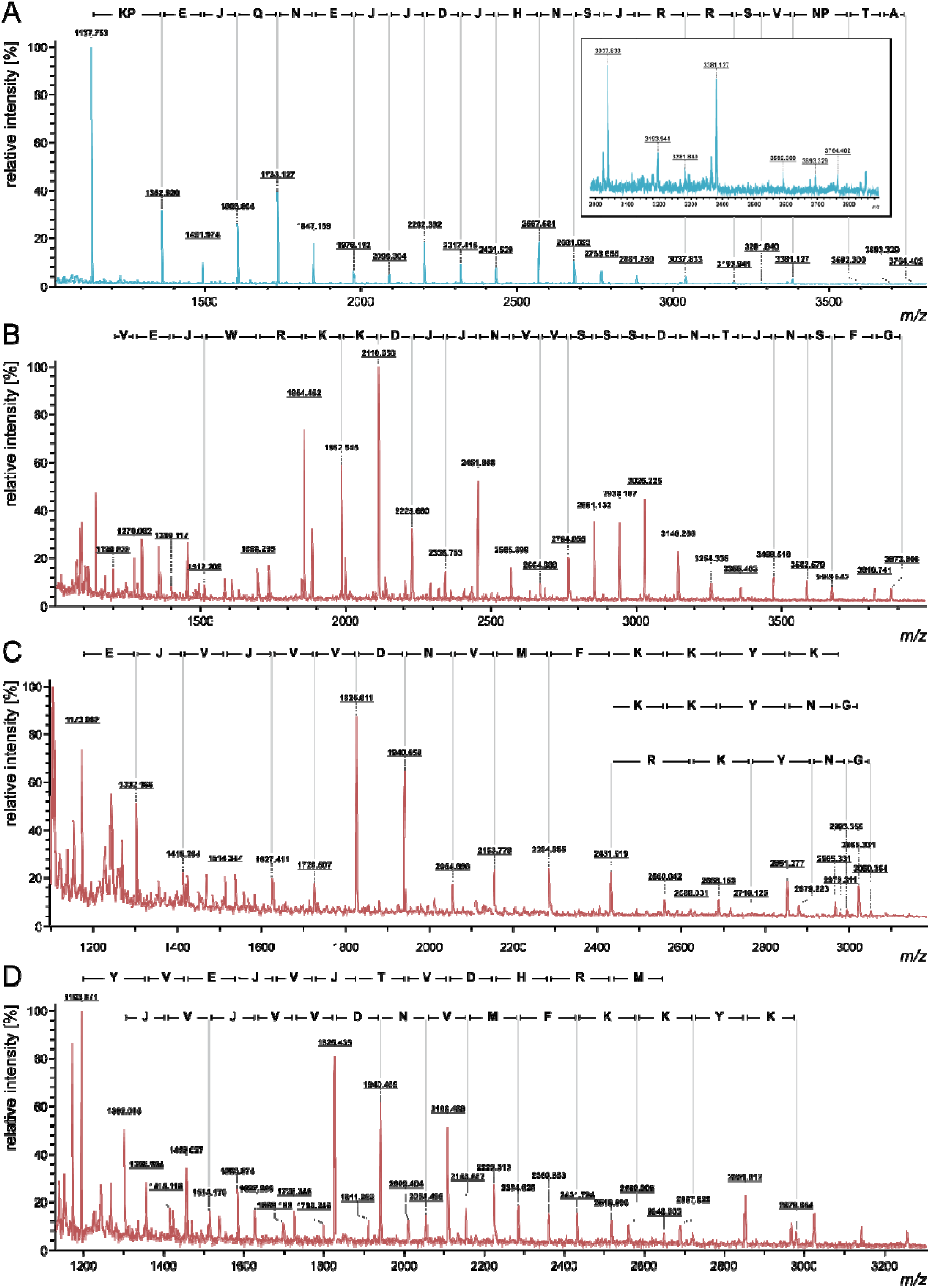
MALDI top-down sequencing by in-source decay of venom components from *Vipera a. senliki*. (**A**) Identification of a cysteine-rich venom protein (CRISP) by N-terminal sequencing (F 16). (**B**) Identification of a snake venom metalloproteinase (svMP) by N-terminal sequencing (F 17). (**C**) Identification of different svMP proteoforms by N-terminal sequencing (F 29). (**D**) Identification of different svMP proteoforms by N-terminal sequencing (F 31/32). No distinction can be made between leucine and isoleucine (J = Leu or Ile).

Interestingly, the majority of top-down ISD sequences are venom proteins of higher molecular weight, e.g. CRISP, svSP and svMP. However, top-down sequencing by ISD-MALDI suffers from low sensitivity and the restriction to highly concentrated, pure protein samples. The presence of highly blended peptide or protein samples (**SI Figure 4**) in tandem MS can lead to an overlap of several *c_n_*-fragment ion series of different protein families, which in turn could be problematic for peptide and protein identification especially in combination with low abundant signals.^73^

### 3.4. Inter-specific venom variation in closely related Eurasian vipers

*Vipera anatolica senliki* is a newly discovered subspecies of *V. anatolica*.^41^ After the description of a new subspecies (*V. a. senliki*), the populations in Kohu Mountain, Çığlıkara Cedar Reserve became nominotypic (*V. a. anatolica*). Therefore, a comparative analysis of venoms between these two subspecies is warranted, and this, in the context of venoms of other closely related and geographically proximal *Vipera* species, could provide insights into inter-specific and geographic venom variation in this region.

The venoms of the two *V. anatolica* subspecies show high similarity, both in peak shape as well as fraction intensities in RP-HPLC chromatograms (**SI-Figure 6**). While the *V. a. anatolica* venom profile is sourced from a previous study, both subspecies were analyzed on the same device with identical quantities of crude venom. The protein families coincide in a direct retention time (t_R_) lineup and by several IMP masses in both venoms (**SI-Table 2**).^18^ Particularly the dominant fractions F 3 (svMP-i), F 4 (BPP), F 9 (DI), F 14 (PLA_2_) and F 16 (CRISP) consist of the same main components. This shows that in the two subspecies, venoms have a similar composition in these toxin families. An exception is disintegrin (DI) subunit dimerization (**Figure 2**, **SI-Figure 5**): While the *V. a. anatolica* venom contains only two dimeric DI (13982.8 Da; 14001.8 Da), the *V. a. senliki* venom includes a third abundant DI mass of 13999.8 Da. The lack of this dominant fraction 10 and correspondingly of the mass in the *V. a. anatolica* venom suggests that the 7022.0 Da subunit is a DI specific to *V. a. senliki*.

An overall comparative analysis of venom from the two subspecies shows similarity in toxin compositions with some particular features (**Figure 5**, **SI-Table 2**). Several svMP variants form the most abundant toxin family in both *V. a. senliki* and *V. a. anatolica* with 42.9% and 41.5%, respectively. This is followed by CRISP, which was at 9.8% in *V. a. senliki* but slightly more dominant in *V. a. anatolica* (15.9%). On the other hand, some venom components are more abundant in *V. a. senliki* as opposed to *V. a. anatolica*, including svSP (7.1% to 1.6%), CTL (4.6% to 1.1%) and Kunitz-inhibitors (1.2% to 0.3%). In contrast, PLA_2_ (∼8%), DI (∼2%) and peptides (∼22%) do not show major variations. It is interesting to note that the toxin family LAAO was found only in *V. a. senliki*. Furthermore, it is worth mentioning, that in *V. a. anatolica* 10% of venom proteins belong to the unknown category and could not be annotated.^18^

**Figure 5:**
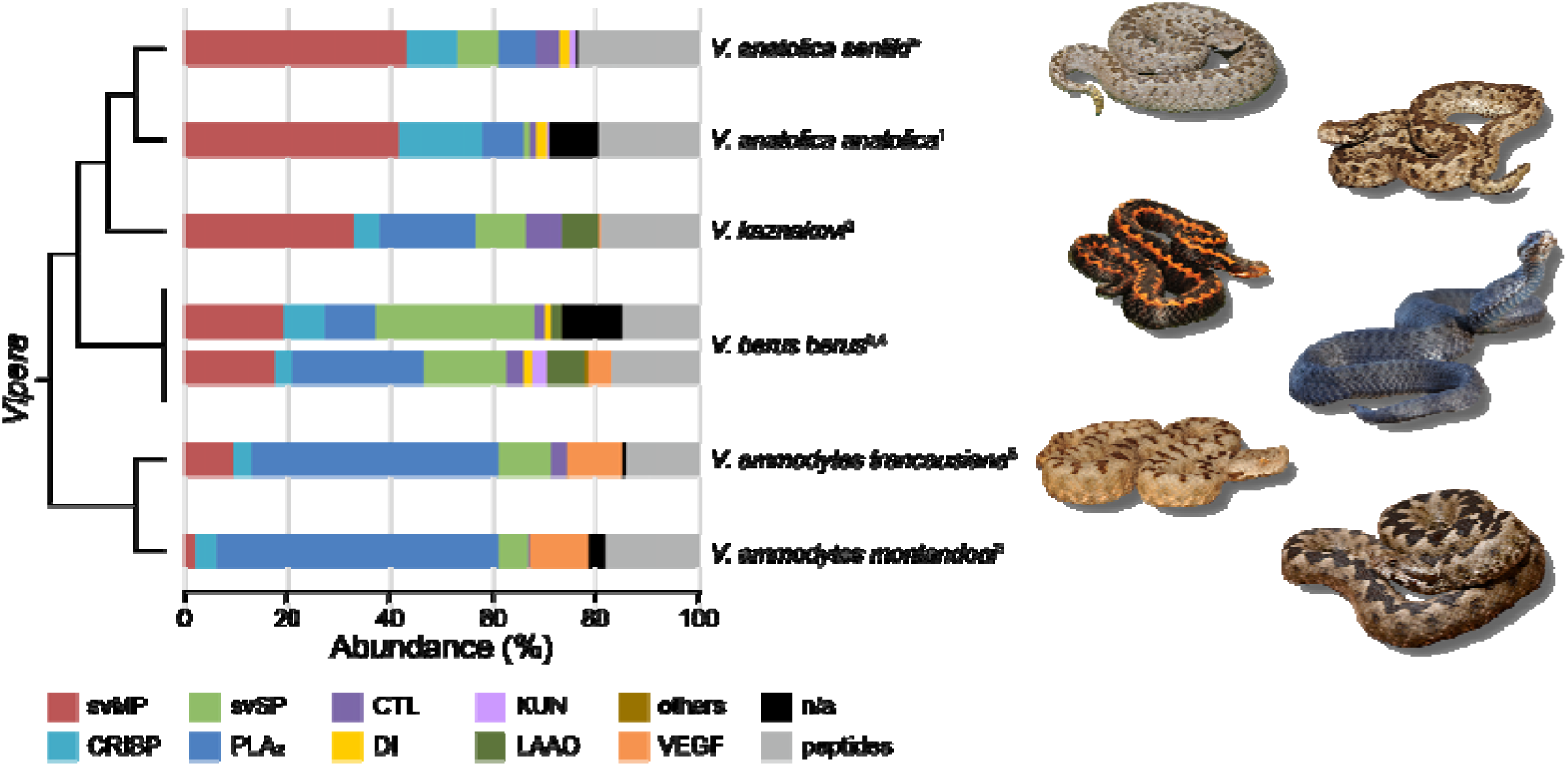
Comparative venom proteomic data from six closely related members of the *Vipera* genus in the region of the Black Sea. Seven venoms of six different *Vipera* (*V*.) species and subspecies are compared by their composition. Taxonomic relations based on Alencar *et al.*^79^ and Göçmen et al.^41^ are shown without genetic distances. The asterisked data set is a result of this study. The origins of toxin ratios are marked top to bottom numerically: 1 - Göçmen *et al.*^18^, 2 - Petras *et al.*^74^, 3 - Latinović *et al.*^75^, 4 - Al-Shekhadat et al.^76^, 5 - Hempel *et al.*^20^. Snake images correspond in the order to the cladogram species.

Further, we reviewed and compared the previously published venomics data of four closely related and geographically proximal *Vipera* species in relation to *V. anatolica* (**Figure 5**). These include the Caucasian *V. kaznakovi*^74^, the Eurasian *V. berus berus*^75, 76^, and two subspecies of *V. ammodytes* occurring in Greece and Turkey, *V. a. montandoni* and *V. a. transcaucasiana*.^20^ All of these studies used a consistent snake venomics workflow: quantification with RP-HPLC, 1D-PAGE and single band bottom-up. While additional quantified venomic studies are available for *V. kaznakovi* (shotgun bottom-up) and *V. berus berus* (2D-PAGE), these were not considered due to variant quantification methods and the direct compositional lineup.^77, 78^ Generally, svMP/DI, CRISP, PLA_2_ and svSP are found in venoms of all species and subspecies, however compositional differences appear to exist between species/subspecies and clades (**Figure 5**). svMPs are the most dominant venom components in the clade containing *V. kaznakovi* and *V. anatolica* subspecies, whereas PLA_2_ is the major component in *V. ammodytes*. The residual toxins are only low abundant in some venoms or vary strongly in their presence, e.g. VEGF which is with 10% most dominant in *V. ammodytes*. A consistent aspect among all species however is the presence of ∼15-20% peptides in the venoms of all *Vipera* venoms compared here. In the case of *V. berus berus*, the two studies show differences in venom compositions, and may reflect intraspecific variation at the population level in large geographic areas of species distribution.^75, 76^ However, this cannot be ascertained as the original localities from which snakes were collected are not available in all cases.

## 4. Conclusions

Herein we describe the detailed venom composition of the newly discovered subspecies *Vipera anatolica senliki* by an integrative combination of venom gland transcriptomics as well as decomplexing bottom-up and top-down proteomics.

The transcriptome sequencing revealed 14 full length toxin family transcripts and 28 partially assembled toxins. The toxin gene expression in *V. a. senliki* is atypical compared to gene expression patterns reported so far for other *Viperinae*. Relative quantitative snake venomics showed nine venom toxin families with snake venom metalloproteinases (svMP, 42.9%) as most abundant protein family, followed by CRISP (9.9%), PLA_2_ (8.2%), svSP (7.2%) and CTL (4.6%) as well as DI (1.9%), KUN (1.2%), LAAO (0.1%) and non-annotated components (0.5%). The comparison of native and chemically reduced venom components by established top-down venomics allowed the identification of three venom protein families by inter- and intramolecular disulfide bonds. Furthermore, we identified a high content of peptides (23.5%), including svMP-inhibitor (svMP-i, 5.9%) and bradykinin potentiating peptides (BPP; 0.6%). The comparative venom analysis of the two *Vipera anatolica* subspecies showed a high similarity in toxin compositions with some particular features. A broader inter-species analysis of closely related viperid venoms, using consistent snake venomics workflow quantification, gave further insights in the venom proteome distribution. Whether the venom compositions of the reviewed species result from phylogenetic relationships, geographic distribution, or dietary variation could be interesting to investigate in future.

In addition, we assessed an in-source decay top-down protocol which also enabled disulfide bond mapping for KUN and PLA_2_ by direct comparison of reduced and non-reduced spectra. In combination with N-terminal top-down sequencing a precise mass fingerprint for a number of high molecular mass venom proteoforms was enabled. Top-down sequencing by ISD-MALDI has the potential to overcome persisting limits of current mass spectrometric techniques. Its exploitation as an alternative sequencing method may help to better characterize venom proteomes in combination with established snake venomics approaches. We showed that ISD top-down sequencing renders a sufficient number of mass fingerprints for the characterization of high molecular mass venom components, typically over-represent in viperid venoms. Moreover, the method allows sufficient mass fingerprints for several proteoforms even present in one chromatographic fraction. Due to the requirements for highly concentrated and pure samples however, it is still a long way for ISD top-down sequencing being a high-throughput workflow. In this context, a second-dimension chromatographic separation technique (e.g. SEC, IEC or EBA) could help to overcome poor chromatographic resolution. The software-assisted spectral assignment of venom proteoforms and related PTMs by top-down ISD spectra is a further limitation due to the lack of reliable bioinformatic tools. The manual assignment of such superimposed fragmentation spectra can be a time-consuming process. Therefore, the development of open-source and user-friendly bioinformatic tools would be a first step to turn ISD top-down sequencing into a rapid assignment tool for a broader community. These aspects are currently the main obstacles that prevent ISD top-down sequencing from being an alternative, rapid terminal sequencing method so far. Nevertheless, due to many advantages ISD top-down sequencing can most likely gain momentum to analyze venom proteomes in its entirety and become a rapid sequencing tool by eliminating existing problems by further experiments and optimizations.

## Author contributions

B.-F.H., M.D. and R.D.S. planned the study. A.N., B.G. and M.K. collected the animals and prepared venom and venom gland tissue samples. B.-F.H., M.D. performed the protein separation and acquired the mass spectrometry data. M. performed the transcriptome sequencing and bioinformatics. B.-F.H., M.D., and M. performed the data analysis. R.M.K. provided infrastructure and resolved toxin sequences. A.N., M. and R.D.S. acquired funding and provided materials and instruments for the study. B.-F.H., M.D. and M. wrote the manuscript with assistance from R.D.S. and R.M.K. All authors read, discussed and approved the manuscript.

## Acknowledgement

We dedicate this paper to the memory of Professor Bayram Göçmen, who lost his fight against cancer. He was an outstanding teacher, a good friend and colleague and loved by his family. Further, we would like to thank Christoph Weise from the Freie Universität Berlin for the permission and access to the MALDI device as well as showing us the basic principles for ISD-MALDI. We also want to thank Joshua Baal for providing us with pictures of *Vipera berus berus* and Bayram Göçmen for all remaining pictures in this study. We thank Murat Şenlik (subspecies name is derived from his surname) and M. Anıl Oğuz for their assistance during the fieldwork.

The transcriptome analysis was founded by a South-East Asian Biodiversity Genomics (SEABIG) Grant No. R-154-000-648-646.

## Conflict of Interest

The authors declare no conflict of interest.

