## Supplementary Information for "Extended snake venomics by top-down in-source decay: Investigating the newly discovered Anatolian Meadow viper subspecies, *Vipera anatolica senliki*"

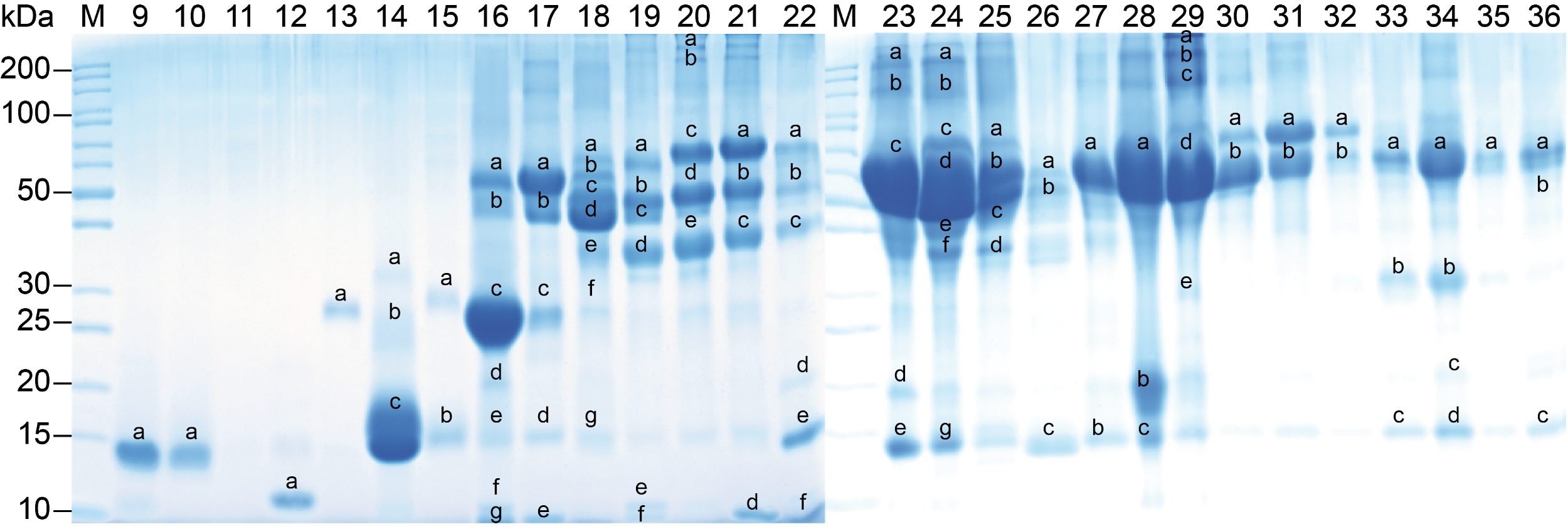

**SI-Figure 1: SDS-PAGE fractions of *Vipera anatolica senliki* venom under reducing conditions.** RP-HPLC venom fractions shown in Figure 1 were further processed by SDS-PAGE analysis. Fraction numbers are indicated above the lanes. Nomenclature shows selected bands for tryptic in-gel digestion and subsequent bottom-up venomics.

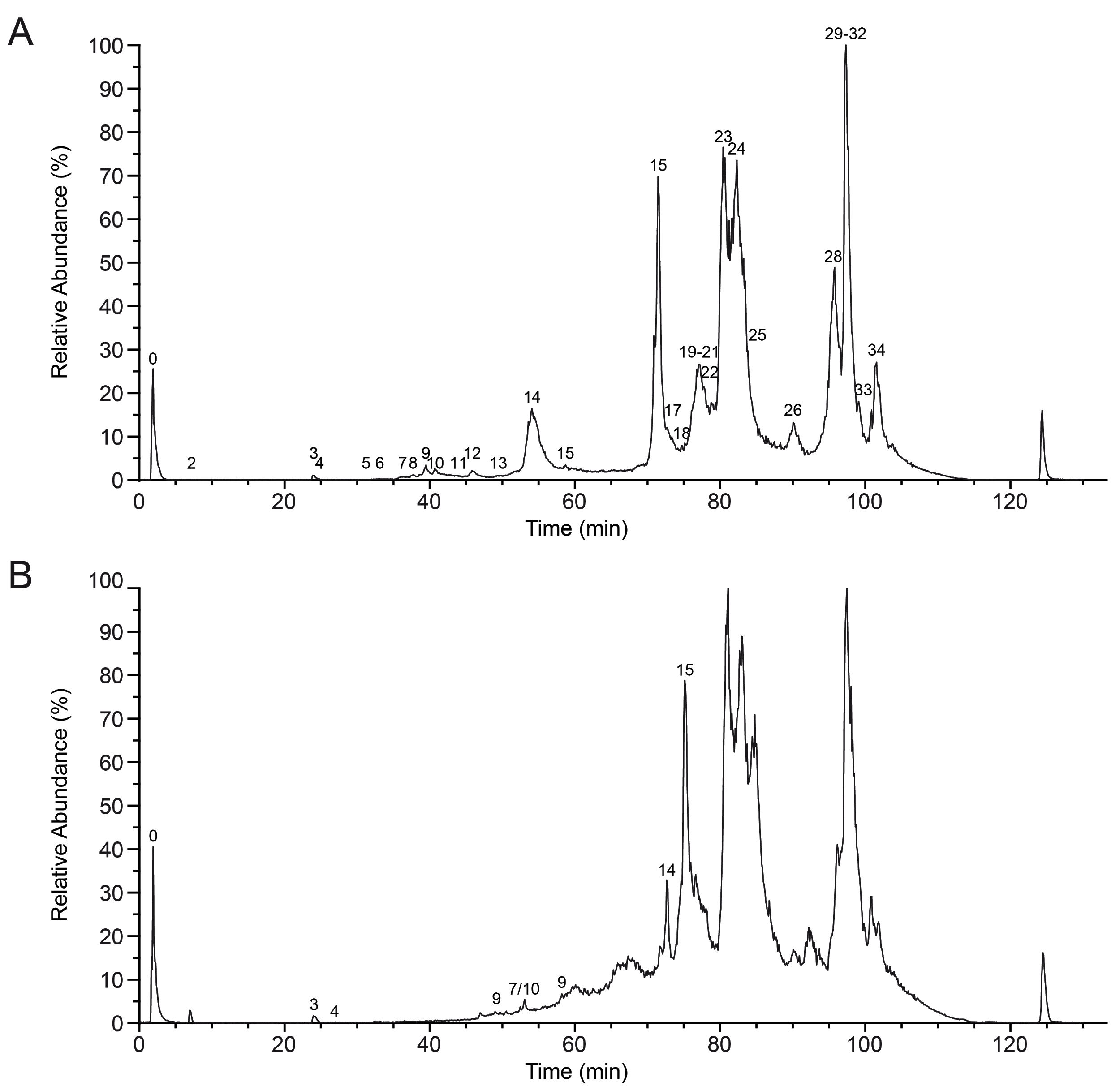

**SI-Figure 2: Extended snake venomic analysis of native and chemically reduced *Vipera anatolica senliki* crude venom.** Total ion chromatogram (TIC) from (**A**) native and (**B**) reduced *V. a. senliki* venom for IMP. The total ion counts were measured by HPLC-ESI-MS and the relative abundance was set to 100% for the highest peak. Fraction nomenclature based on Figure 1.

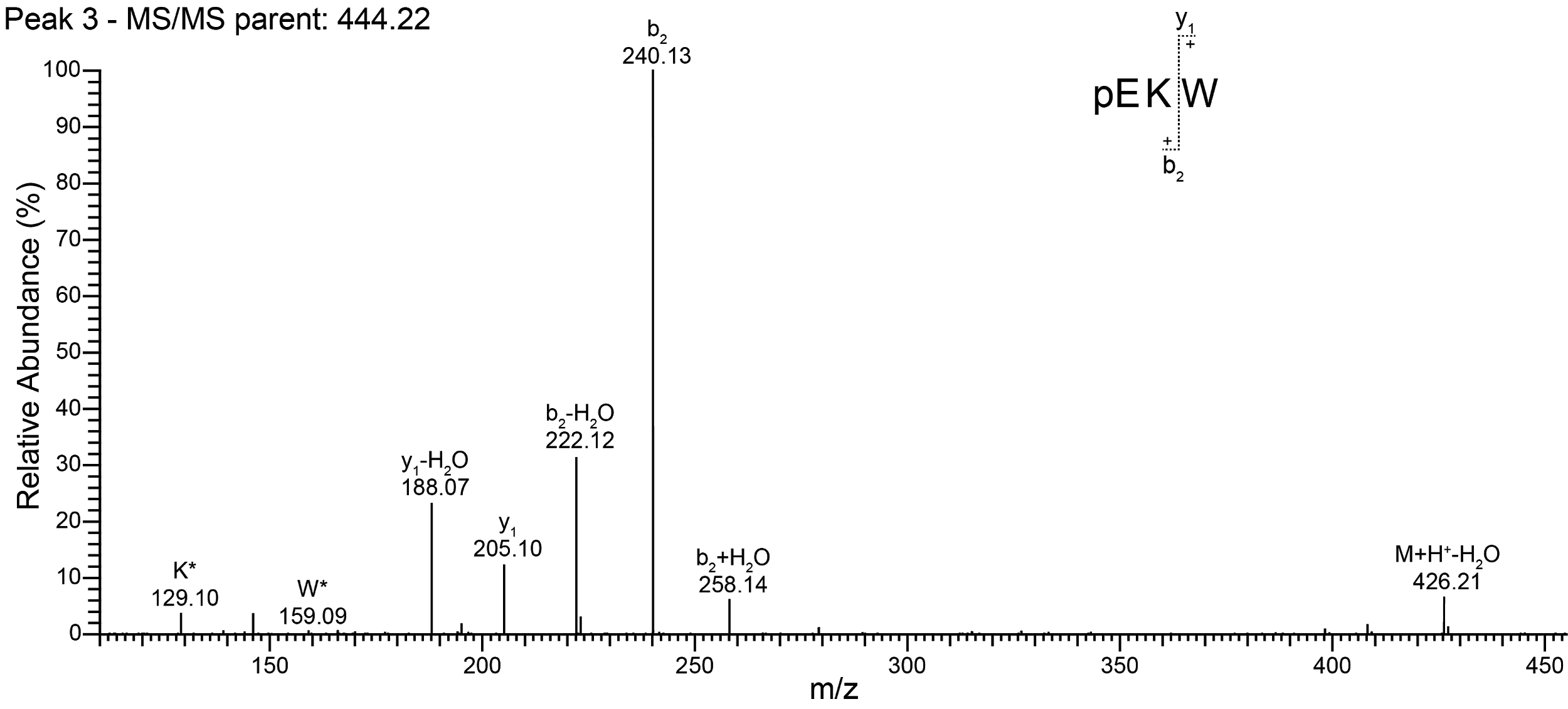
**SI-Figure 3: Tandem MS spectrum of the tripeptidic metalloprotease inhibitor pEKW.** Representative MS/MS spectra of a small tripeptic svMP inhibitor (svMP-i) with *m/z* 444.22 precursor ion mass for *de novo* annotation in the *Vipera anatolica senliki* venom.

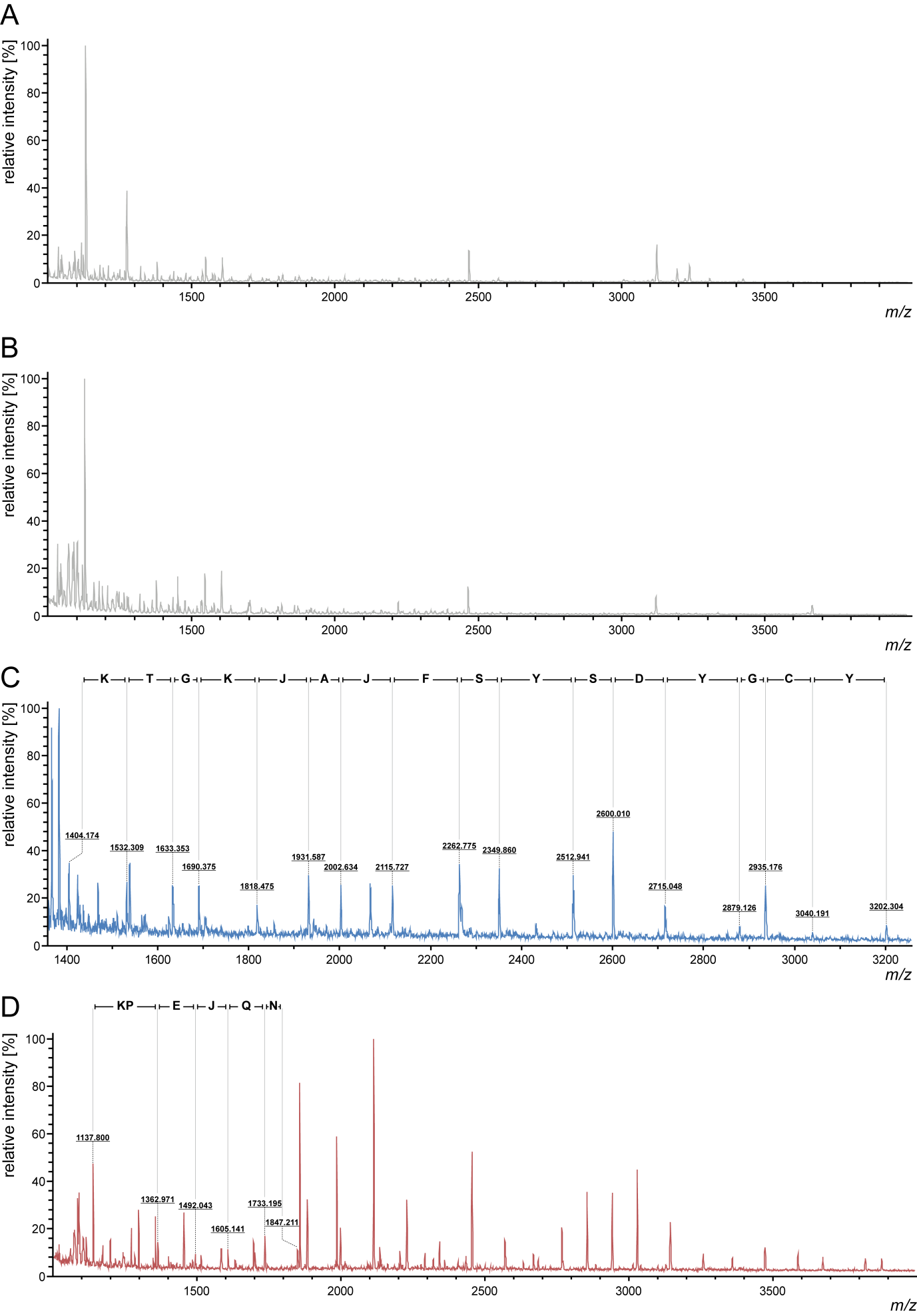

**SI-Figure 4: MALDI top-down sequencing by in-source decay of different venom components from *Vipera a. senliki*.** (**A**) Examples of top-down ISD spectra of peptide fractions (F 8) and (**B**) (F 9/10) showing no distinct sequences. (**C**) Identification of a phospholipase A_2_ (PLA_2_) proteoform (ammodytin I2 (D)) by N-terminal sequence (F 14/15). (**D**) Identification of a short cysteine-rich venom protein (CRISP) peptide fragment by N-terminal sequencing (F 17), previously annotated in peak 16. No distinction can be made between leucine and isoleucine (J = Leu or Ile).

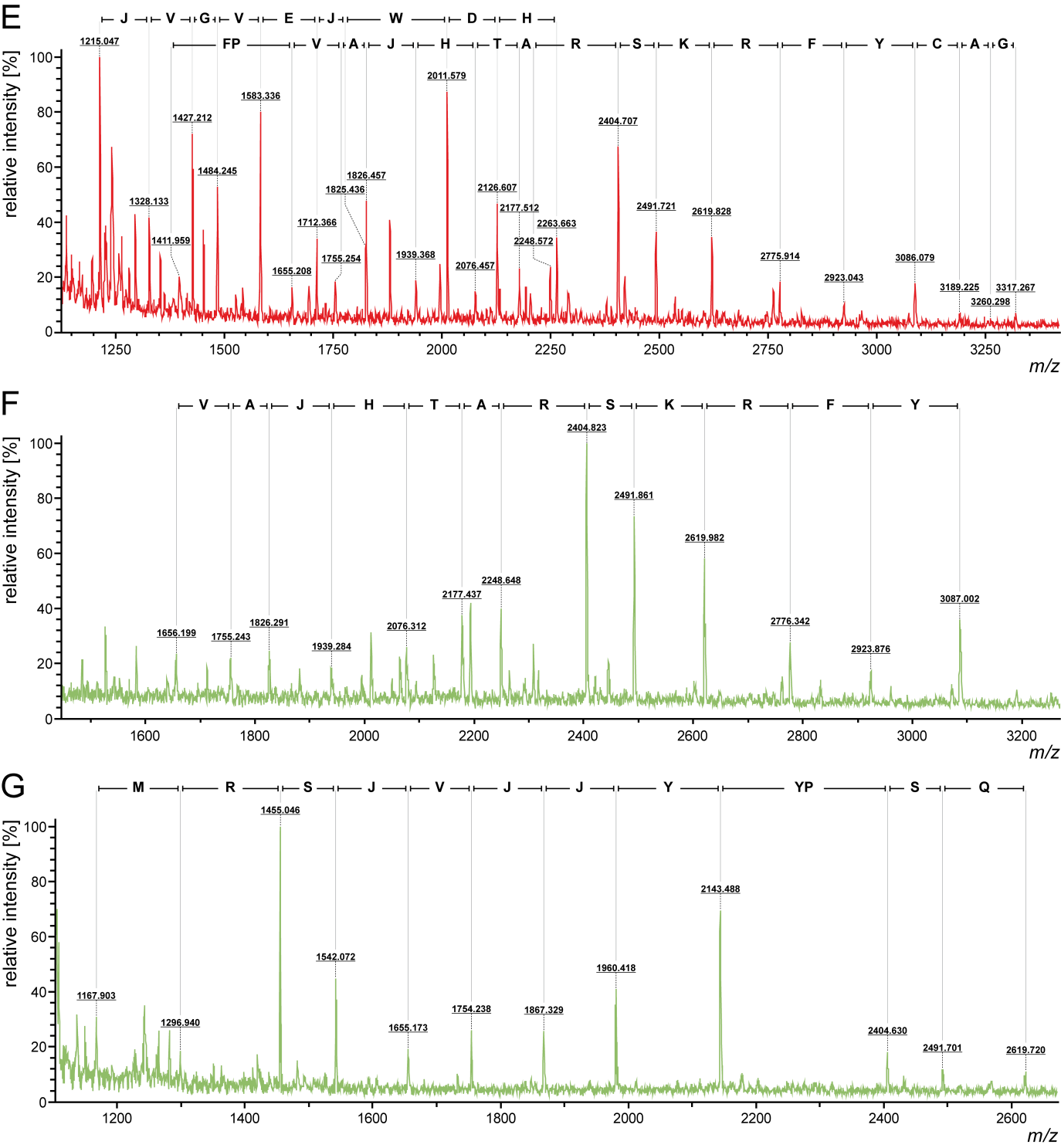

**SI-Figure 4 *(continued)*: MALDI top-down sequencing by in-source decay of different venom components from *Vipera a. senliki*.** (**E**) Identification of a snake venom metalloproteinase (svMP) proteoform by N-terminal sequencing (F 18-22). (**F**) Identification of a snake venom serine protease (svSP) proteoform by N-terminal sequencing (F 23) with a specific transcriptome hit (VT_T0953_R_0_0019_L_1199_SP). (**G**) Identification of a svSP proteoform by N-terminal sequencing (F 24). No distinction can be made between leucine and isoleucine (J = Leu or Ile).

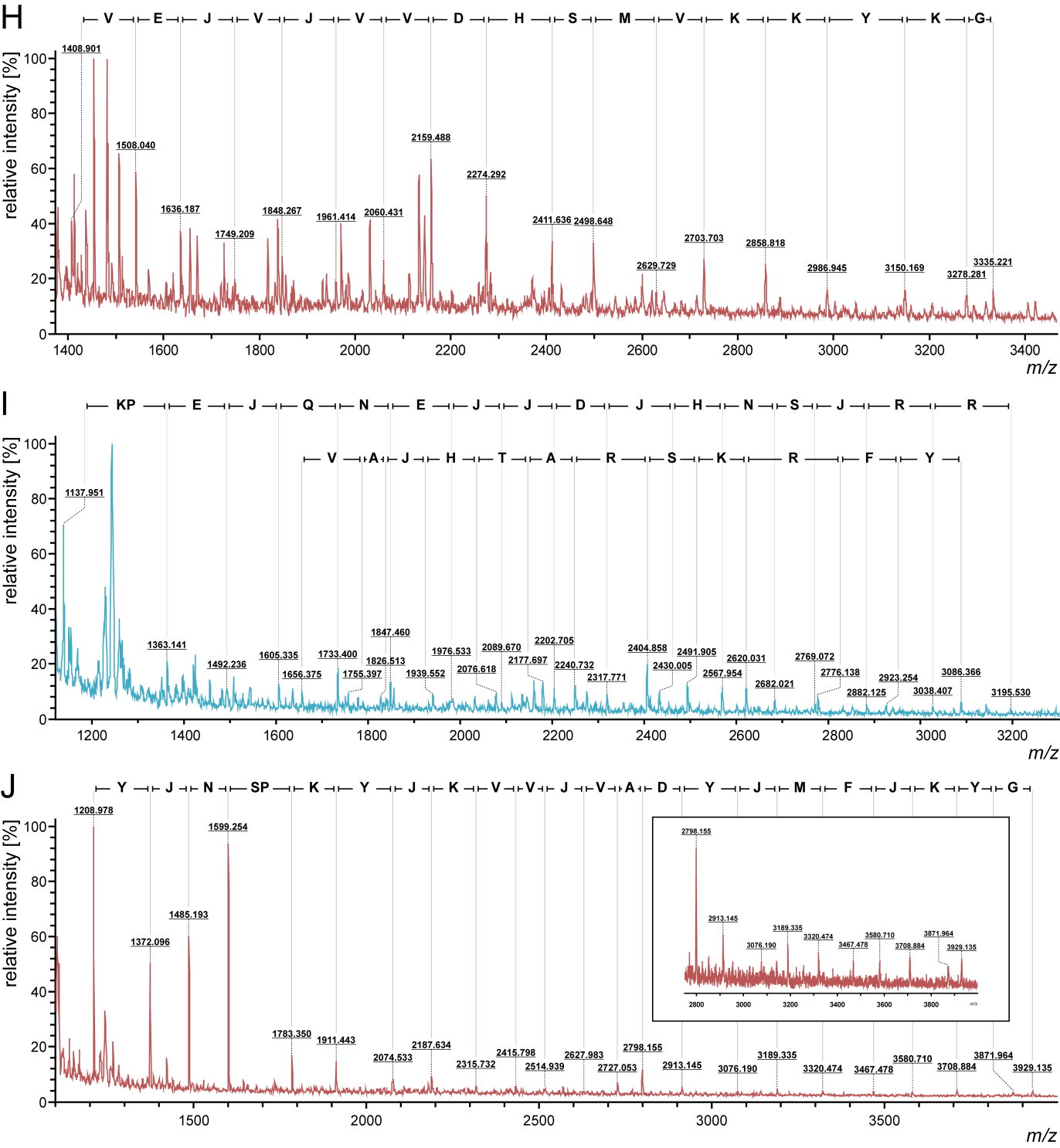

**SI-Figure 4 *(continued)*: MALDI top-down sequencing by in-source decay of different venom components from *Vipera a. senliki*.** (**H**) Identification of a svMP proteoform by N-terminal sequencing (F 25) with a specific transcriptome hit (DN2248_c0_g1_i1_len_747_SVMP). (**I**) Identification of a CRISP proteoform with a specific transcriptome hit (DN8323_c0_g1_i1_len_755_CRISP) and a svSP proteoform with a specific database hit (VT_T0953_R_0_0019_L_1199_SP) (F 27). (**J**) Identification of a svMP proteoform by N-terminal sequencing (F 33-35). No distinction can be made between leucine and isoleucine (J = Leu or Ile).

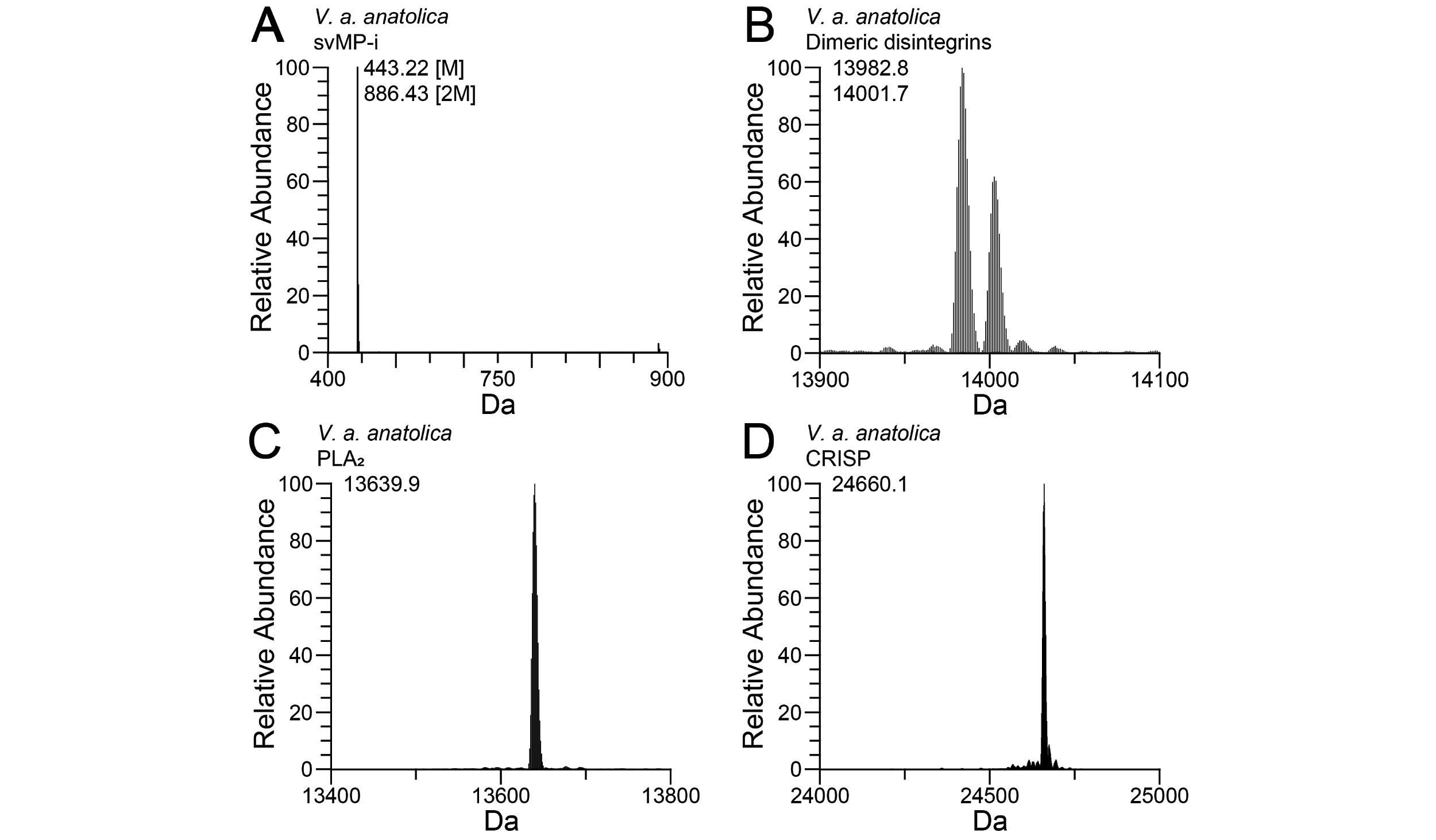

**SI-Figure 5: Intact mass profiling of exemplary *Vipera anatolica anatolica* venom components.** *V. a. anatolica* (Göcmen *et al.*^1^) shows compared to *V. a. senliki* (**Figure 2** and **SI-Table 2**) identical toxin masses, like (**A**) svMPI-i, (**B**) dimeric disintegrins, (**C**) PLA_2_ and (**D**) CRISP.

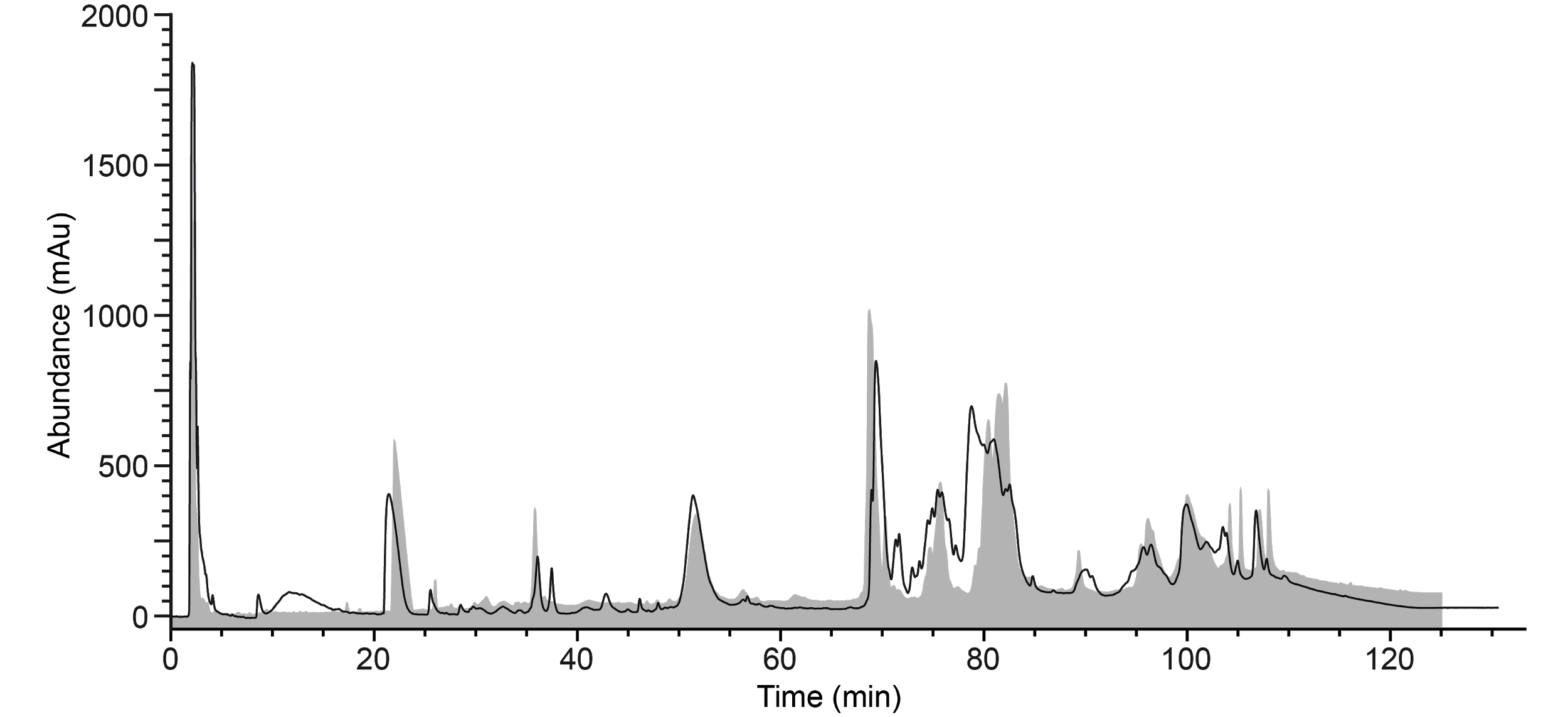

**SI-Figure 6: Overlay of C18 RP-HPLC venom profile from *Vipera anatolica senliki* and *Vipera anatolica anatolica*.** HPLC venom profile of *V. a. senliki* (black line) is shown compared to the *V. a. anatolica* (grey) analysis by Göçmen *et al.*^1^. Same venom amounts of venom were measured on identical devices and column.

**Table S1. Venom proteins and peptides identified from *Vipera anatolica senliki.*** Assignments of venomic components by crude venom intact mass profiling (IMP, method A), IMP of a single RP-HPLC fraction with low molecular mass (method B), bottom-up (BU, method C) and in-source decay annotation (ISD, method D). Fraction numbers are based on the RP-HPLC chromatogram (**Figure 1**). Annotation was performed *de novo* and by peptide spectrum matching from in-gel digested protein bands (**SI-Figure 1**). Identification was carried out against a non-redundant *Viperidae* protein database (taxid: 8689), our custom transcriptome database and a set of proteins found as common contaminants (cRAP). SDS-PAGE and intact mass profile analysis provided the average molecular weight. Most abundant mass for IMP analysis s is marked by *. IMP performed by charge-state deconvolution was carried out with MagicTransformer (MagTran) and is marked by #. The green-marked entries are peptide sequences found by our ISD top-down approach.

**SI-Table 2. Compositional venom lineup of two *Vipera anatolica* subspecies.** The most abundant toxin families in the venoms of *Vipera anatolica senliki* (this study) and *Vipera anatolica anatolica* (Göçmen *et al.*^1^) are compared by their HPLC retention time. Venoms were measured on identical devices and column. Identical identified masses in the IMP are mentioned in the correspondent row.

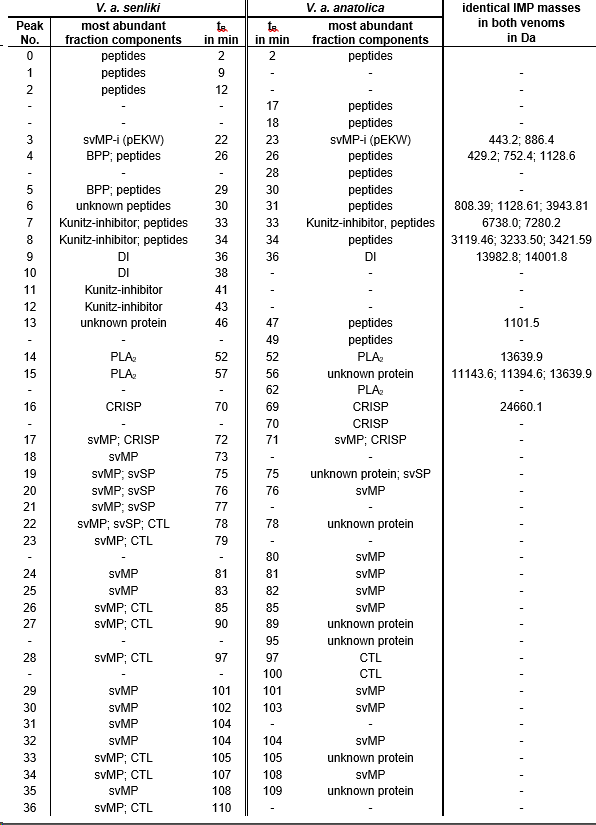
